# Transcription-coupled and epigenome-encoded mechanisms direct H3K4 methylation

**DOI:** 10.1101/2021.06.03.446702

**Authors:** Satoyo Oya, Mayumi Takahashi, Kazuya Takashima, Tetsuji Kakutani, Soichi Inagaki

## Abstract

Mono-, di-, and trimethylation of histone H3 lysine 4 (H3K4me1/2/3) are associated with transcription, yet it remains controversial whether H3K4me1/2/3 promote or result from transcription. Our previous characterizations of *Arabidopsis* H3K4 demethylases suggest roles for H3K4me1 in transcription. However, the control of H3K4me1 remains unexplored in *Arabidopsis*, in which no methylase for H3K4me1 has been identified. Here, we identified three Arabidopsis methylases that direct H3K4me1. Analyses of their genome-wide localization using ChIP-seq and machine learning revealed that one of the enzymes cooperates with the transcription machinery, while the other two are associated with specific histone modifications and DNA sequences. Importantly, these two types of localization patterns are also found for the other H3K4 methylases in *Arabidopsis* and mice. These results suggest that H3K4me1/2/3 are established and maintained via interplay with transcription as well as inputs from other chromatin features, presumably enabling elaborate gene control.

## Introduction

Posttranslational modifications of histone residues shape the epigenome and affect gene expression ^1^. Histone H3 lysine 4 methylation (H3K4me) is a conserved histone mark found in actively transcribed genes; however, the molecular basis and functional significance of the interrelationship between H3K4me and transcription remain elusive ^2, 3^. It has been proposed that H3K4me can be viewed as a memory of transcription because transcription directs H3K4 methylation ^4–7^. On the other hand, in some experimental systems, H3K4me is deposited independent of transcription ^8, 9^. The underlying mechanisms that coordinate transcription-coupled and transcription-independent H3K4me are largely unexplored.

Another layer of H3K4me complexity is its occurrence in three states (mono-, di-, and trimethylation; me1, me2 and me3, respectively). Generally, H3K4me3 and H3K4me2 are found on the first several nucleosomes of genes, while H3K4me1 is found in downstream regions within the gene bodies ^10–12^. Our previous genetic and genomic studies in *Arabidopsis thaliana* (hereafter *Arabidopsis*) have demonstrated key roles of gene body H3K4me1 in the control of transcription; a putative H3K4me1 demethylase, LYSINE-SPECIFIC DEMETHYLASE1-LIKE2 (LDL2), mediates transcriptional silencing induced by gene body H3K9me2 ^13^, while a related demethylase, FLOWERING LOCUS D (FLD), coordinates convergent overlapping transcription by removing H3K4me1 ^14^. Although these findings regarding the H3K4 demethylases of *Arabidopsis* suggest a link of H3K4me1 with other chromatin modifications and transcription, the mechanisms of H3K4me1 control remain elusive, mainly because methylase(s) for H3K4me1 has not been identified in plants.

In both animals and plants, two types of H3K4 methylases are known: Set1-type and Trithorax/Trithorax-related (Trx/Trr)-type H3K4 methylases ^15^. However, yeasts have lost Trx/Trr-type H3K4 methylases during evolution ^16^, and yeast Set1 is the sole H3K4 methylase responsible for all H3K4me1/2/3. *Arabidopsis* has five Trx/Trr-type enzymes (Arabidopsis Trithorax (ATX) 1 to 5) and at least one Set1-type enzyme (Arabidopsis Trithorax Related (ATXR) 7) (Fig. 1a). Additionally, ATXR3, which has a domain structure characteristic of Set1-type enzymes but contains an atypical catalytic domain as H3K4 methylase ^17, 18^, can catalyze all three states of H3K4me1-3 *in vitro* ^19^. Loss-of-function mutations in the *ATXR3* gene cause substantial H3K4me3 loss genome-wide, but H3K4me1 and H3K4me2 are largely unaffected ^19, 20^. In addition, triple loss-of-function of *ATX3*, *ATX4*, and *ATX5* leads to global decreases in H3K4me3 and H3K4me2 but not H3K4me1 ^21^. However, methylase mutants with decreased H3K4me1 levels have not been reported. The functions of *ATX1*, *ATX2*, and *ATXR7* genes have been characterized in the context of development, flowering regulation, and plant immunity ^19–27^. For example, they were shown to redundantly inhibit flowering by activating the transcription of the flowering repressor *FLOWERING LOCUS C* (*FLC*) by increasing H3K4me1/2/3 levels within the *FLC* locus ^22, 25^. In the control of *FLC*, ATXR7 was shown to counteract FLD ^25^, which demethylates H3K4me1^1^^4^. However, the genome-wide impacts of these putative H3K4 methylases have not been elucidated.

**Fig.1.**
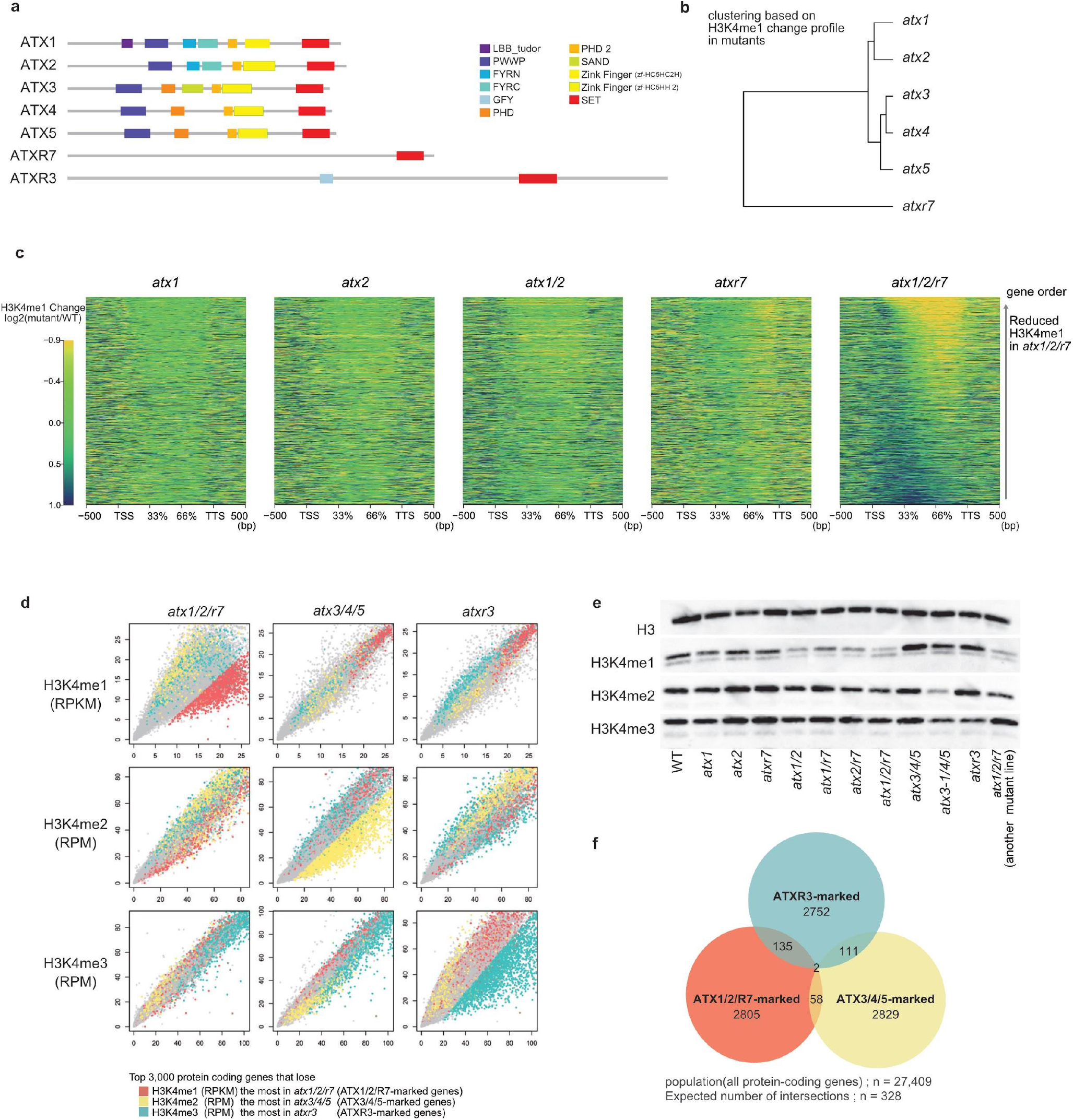
*ATX1, ATX2,* and *ATXR7* redundantly contribute to H3K4me1. **(a)** Domain architectures of ATX(R) proteins from the UniProt database. **(b)** *atx1,2,3,4,5* and *atxr7* mutants were clustered based on the H3K4me1 pattern detected in ChIP-seq (see Methods). **(c)** Mutations of *atx1*, *atx2,* and *atxr7* synergistically cause reduced H3K4me1 in the gene body. H3K4me1 changes from WT are visualized as heat maps. All genes are ordered so that genes that lost H3K4me1 the most in *atx1/2/r7* come to the top. **(d)** ChIP-seq for H3Kme1, H3K4me2, and H3K4me3 in mutants (y-axis) compared to WT (x-axis). Each dot represents each gene. Axes are trimmed to 0.98 quantiles. Top genes (n= 3,000) that lose H3K4me1 most in *atx1/2/r7* (hereafter ATX1/2/R7-marked genes), lose H3K4me2 in *atx3/4/5* (ATX3/4/5-marked genes), and lose H3K4me3 in *atxr3* (ATXR3-marked genes) are colored red, yellow, and blue, respectively. Values for H3K4me2,3 are RPM normalized instead of RPKM as in H3K4me1 because unlike H3K4me1, which covers the gene body, the amounts of H3K4me2 and H3K4me3 are less dependent on gene length ^10^. **(e)** Western blotting of H3K4 methylations on bulk histone extracted from the mutants. **(f)** Activities of ATX1/2/R7, ATX3/4/5, and ATXR3 are observed in mutually exclusive genes. All pairwise ATX1/2/R7-marked genes, ATX3/4/5-marked genes, and ATXR3-marked genes have significantly fewer overlaps than expected (n=328). Significances of the under-representation are tested by hypergeometric tests (p < 1 x 10e-4 for all pairs).

Here, we explored the involvement of seven H3K4 methylase genes in H3K4me1 as well as H3K4me2/3 by analyzing single and multiple mutants using chromatin immunoprecipitation sequencing (ChIP-seq). H3K4me1 levels are substantially decreased by the simultaneous loss of the *ATX1*, *ATX2*, and *ATXR7* genes. Our results clarify the division of labor among H3K4 methylase genes and provide powerful genetic materials for elucidating the functions of each H3K4 methylation state. In addition, we analyzed the genomic localization of these enzymes. The subsequent application of machine learning algorithms revealed that the ATXR7 protein colocalizes with the transcription machinery, while ATX1 and ATX2 are associated with other chromatin modifications (epigenome) and specific DNA sequences (genome). These two types of localization patterns were also found by the reanalysis of other Set1- and Trx/Trr-type H3K4 methylases, including those of animals. These findings lead us to a new perspective: some H3K4me marks may function as records of transcription, while other H3K4me marks may function as mediators of information encoded in the epigenome and/or genome.

## Results

### H3K4me1 levels are decreased by the triple loss of *ATX1/2/R7* genes

To identify enzyme(s) involved in H3K4me1, we examined mutants of six predicted H3K4 methylases: ATX1 to 5 and ATXR7 (Fig. 1a). Among the six single mutants, *atx2* and *atxr7* showed relatively large decreases in H3K4me1 levels (Supplementary Fig. 1a). Interestingly, the *atx2* and *atxr7* mutants differed in the affected regions within genes; *atxr7* showed a marked decrease in H3K4me1 in the 3’ half of the gene bodies, while *atx2* showed a decrease over a broader region (Supplementary Fig. 1b). Cluster analyses based on the intragenic H3K4me1 pattern in the *atx1 to 5* and *atxr7* mutants revealed that *atx1* and *atx2* formed one cluster, while *atx3, atx4, and atx5* formed another cluster (Fig. 1b). These similarities in the effects on the H3K4me1 profile coincide with the similarity of the domain architecture of the corresponding proteins (Fig. 1a).

As the lack of a strong effect in each of the single mutants may reflect redundancy, we examined mutants that lose function of multiple ATX(R) genes. Based on the similarities of the protein domain structures and the effects on H3K4me1 patterns, we examined *atx1/atx2* double mutants and found stronger effects on H3K4me1 than were observed in either of the single mutants (Fig. 1c, Supplementary Fig. 1c). Additionally, *atx1/atxr7* and *atx2/atxr7* showed stronger H3K4me1 decreases than each single mutant (Supplementary Fig. 1c). The effect was still stronger in triple *atx1/atx2/atxr7* mutants (hereafter referred to as *atx1/2/r7*) (Fig. 1c, d, Supplementary Fig. 2a). Western blot analyses confirmed that triple *atx1/2/r7* mutation (with two different sets of T-DNA insertion alleles) resulted in the loss of H3K4me1, while H3K4me2 and H3K4me3 were largely unaffected (Fig. 1e). Thus, ATX1, ATX2, and ATXR7 redundantly contribute to H3K4me1.

Consistent with previous reports ^19–21^, our Western blot and ChIP-seq analyses showed that H3K4me3 and H3K4me2 modifications are mainly mediated by ATXR3 and ATX3/4/5, respectively (Fig. 1d, e; Supplementary Fig. 2b-d). Collectively, the results showed that H3K4me1, H3K4me2, and H3K4me3 levels were specifically reduced in *atx1/2/r7*, *atx3/4/5*, and *atxr3*, respectively (Fig. 1d, e). The genes that were strongly affected (colored red, yellow, or blue in Fig. 1d) in each of these mutants had specific characteristics (Supplementary Fig. 1d) and showed high corresponding H3K4me levels in the wild type (e.g., ATX1/2/R7-marked genes showed high H3K4me1 levels relative to those of other genes) (Supplementary Fig. 1e-g). Interestingly, the target genes of each ATX(R) group were mutually exclusive (Fig. 1f), suggesting that distinct mechanisms direct each of the H3K4 methylases to distinct target genes.

### ATXR7 localization is associated with RNAP2, while ATX1 and ATX2 localization is associated with other chromatin modifications

To investigate the mechanisms that specify the targets of each ATX(R) protein that regulates H3K4me1, we determined the genome-wide localization of the ATX1, ATX2 and ATXR7 proteins using transgenic plants expressing FLAG-tagged proteins (Fig. 2a). ChIP-seq analyses revealed that ATX1 and ATX2 localized around transcription start sites (TSSs), typically in the range of -150 ∼ +300 bp from the TSS. ATXR7 localized around transcription termination sites (TTSs) in the range of -200 ∼ +200 bp from the TTS (Fig. 2b-d). Genes showing H3K4me1 loss in *atx1/2/r7* mutants tended to be bound by ATX1, ATX2 and ATXR7 (Supplementary Fig. 3), supporting the conclusion that these enzymes mediate H3K4me1 on chromatin.

**Fig.2.**
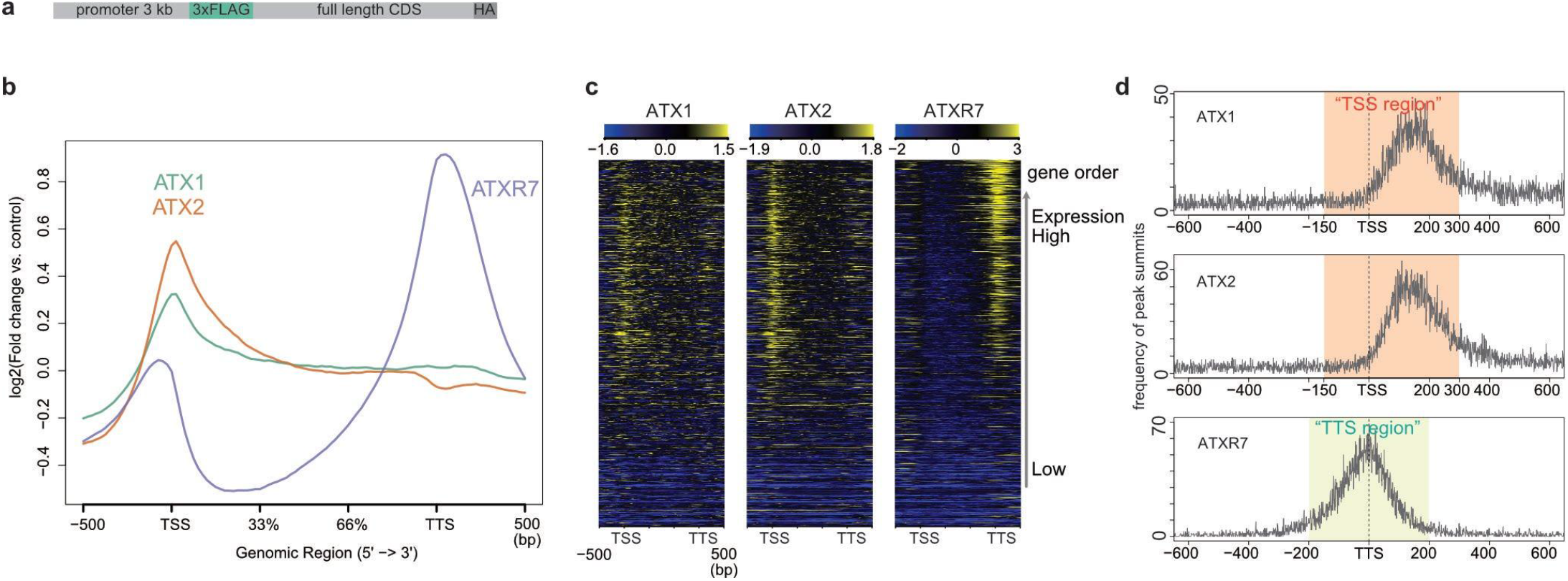
ATX1 and ATX2 occupy TSS while ATXR7 occupies TTS. **(a)** The transgene structure of tagged ATX(R) proteins used for ChIP-seq. **(b)** Metaplot and **(c)** heat maps illustrating ATX1, ATX2, and ATXR7 distributions in gene body region corrected with a non-transgenic control. The heat map was sorted so that highly transcribed genes (measured as mRNA-seq in WT) come to the top. **(d)** Position of ATX1 and ATX2 ChIP-seq peaks relative to TSS and ATXR7 peaks relative to TTS (x-axis), visualized as a frequency of peak summits (y-axis), which are detected against non-transgenic control using MACS2 peak caller. Most of the ATX1 and ATX2 peaks belong to the region spanning from 150 bp upstream to 300 bp downstream of TSS (hereafter ‘TSS region’). Most of the ATXR7 peaks are between 200 bp upstream and 200 bp downstream of TTS (hereafter ‘TTS region’).

To elucidate the chromatin-targeting mechanism(s) of these ATX1/2/R7 proteins, we screened for the determinants of ATX1/2/R7 localization by using a machine learning algorithm (random forest). Using chromatin and genomic features listed in Fig. 3a, the algorithm was trained to distinguish ATX1/2/R7-bound genes from unbound genes. After training, the random forest algorithm reported the ‘importance (mean decrease in Gini)’ of each feature as a relative score reflecting how informative the feature was for classification (Fig. 3a-c). In principle, the values of the features with high ‘importance’ scores can be positively (colocalize) or negatively (exclusive) correlated with the localization level of the corresponding ATX1/2/R7 protein. By taking all the features into account, the localization of each ATX1/2/R7 protein was well explained (Fig. 3d-f; note the high area under the curve (AUC) values).

**Fig.3.**
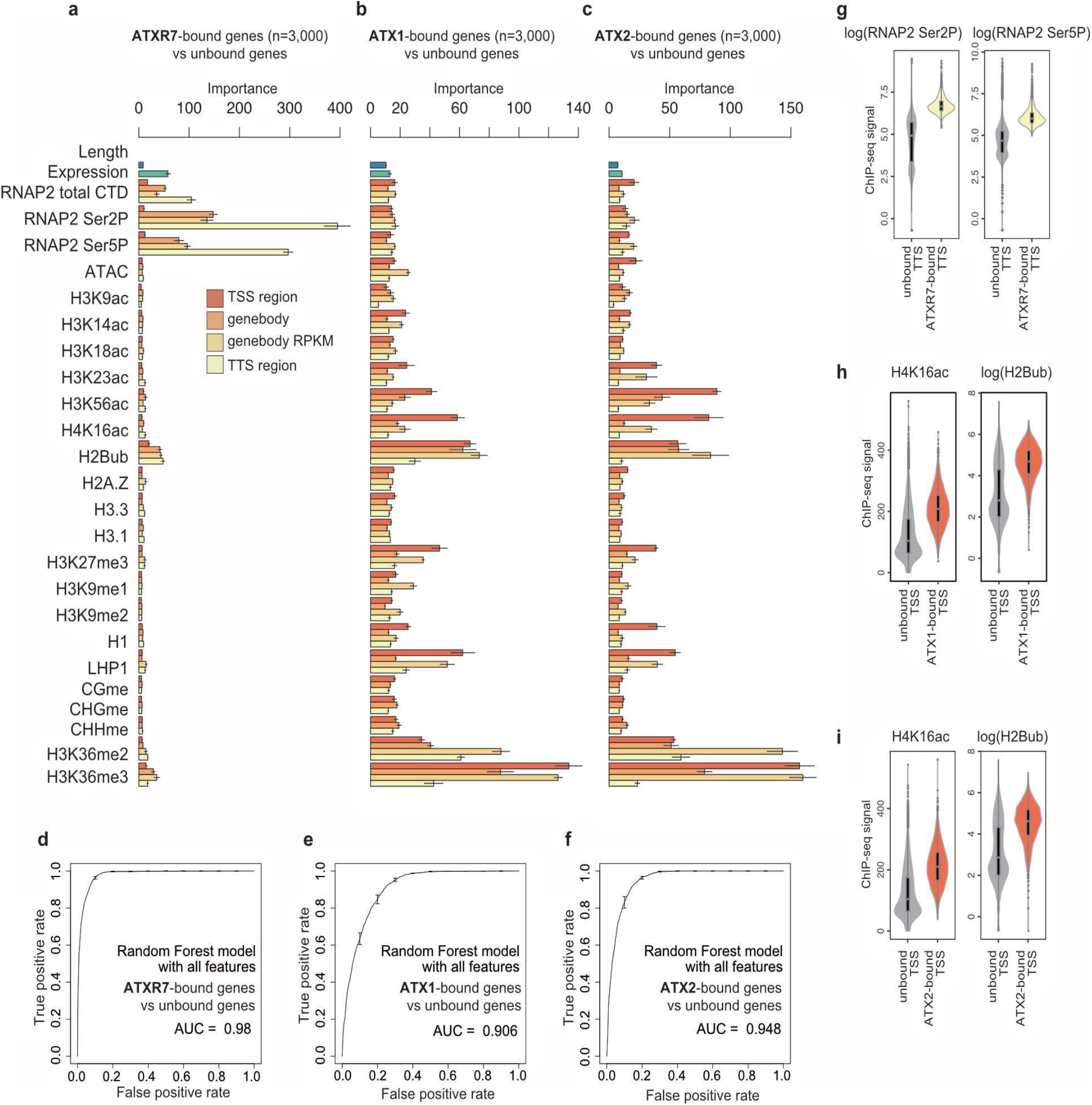
ATXR7 localization is associated with RNAP2, while ATX1 and ATX2 localization is associated with other chromatin modifications. **(a-c)** Chromatin features predictive of ATXR7 (**a**), ATX1 (**b**), and ATX2 (**c**) localization. ATXR7(ATX1,2)-bound genes were defined as the top 3,000 genes with the highest ChIP-seq signal in ‘TTS region (or TSS region)’ compared to nontransgenic control. Random forest models were trained to predict ATXR7(ATX1,2)-bound and unbound genes. Bars indicate ‘mean decrease in Gini’ or in other words ‘importance’ derived from the random forest model, which is a relative score reflecting how informative the feature was for classification. ‘Importances’ are averaged from 5 repeats of training, each of which independently chose negative samples (unbound genes). Error bars are the standard deviation of the 5 repetitions. **(d-f)** ROC plots showing the prediction accuracy of the random forest models. AUC indicates the area under the ROC curve. ROC and AUC are calculated with data on Chr 5, which were held out from training as test data. Average and standard deviation of the 5 repeats of training are plotted. **(g-i)** Violin plots showing the abundance of two most predictively ‘important’ features for each protein. Analyses with ChIP-seq datasets of biological replicates showed similar results (Supplementary Fig. 6).

**Fig. 4.**
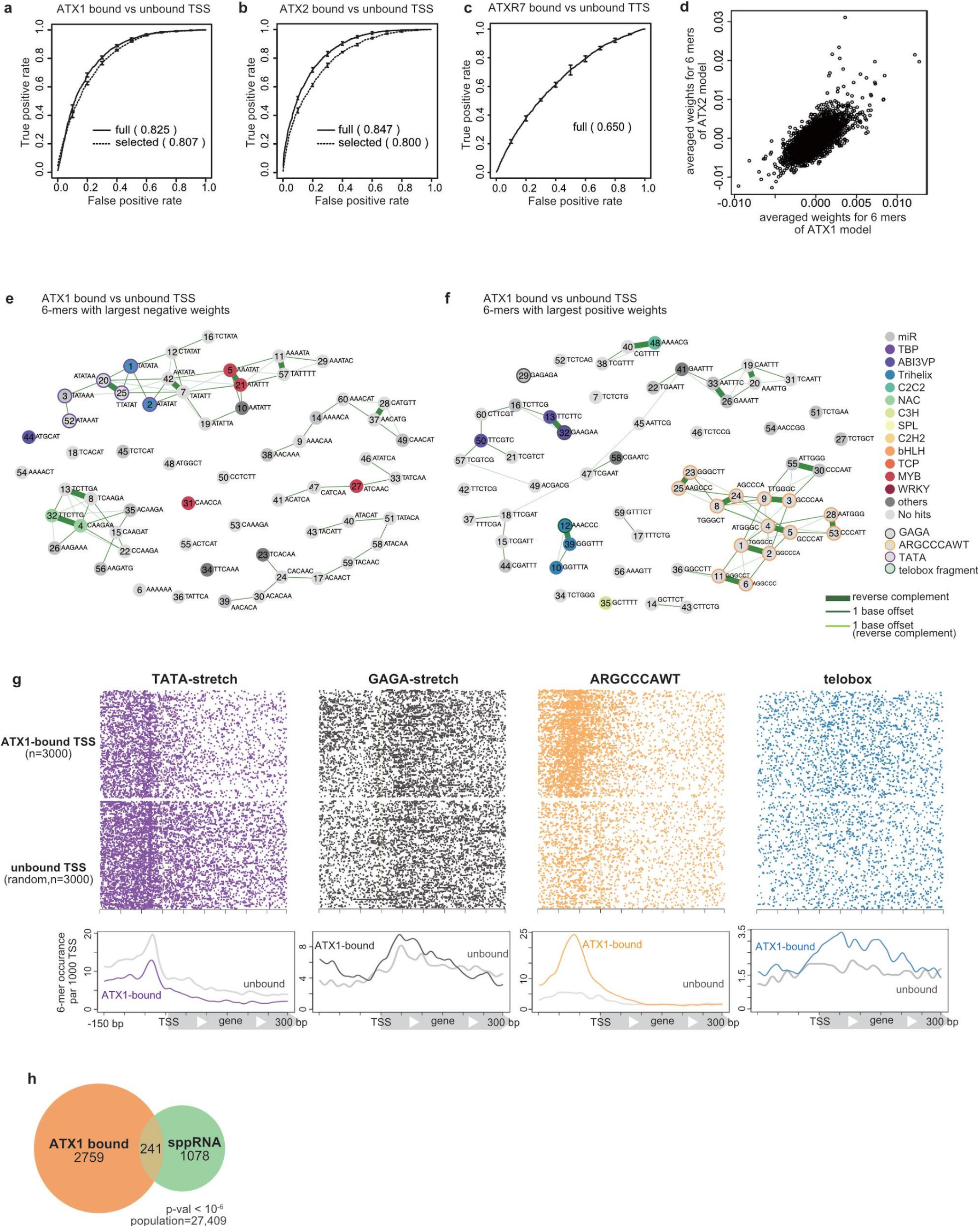
The DNA sequences of ATX1- and ATX2-bound TSSs have distinct architectures. **(a-c)** ROC plots that show predicting accuracy of linear SVM models, which are trained with the full set of 6-mers (solid line) and trained with the 120 6-mers which were highly weighted in the full model (dashed line), to discriminate **(a)** ATX1- or **(b)** ATX2-bound vs -unbound TSS DNA sequence, and **(c)** ATXR7-bound and-unbound TTS sequence. ROC and AUC are calculated with data on Chr 5, which are held out from the training as test data. Averaged scores of 5 cross-fold validation models are plotted. Analyses with ChIP-seq datasets of biological replicates (see Methods) showed similar results (Supplementary Fig. 8) **(d)** Averaged SVM weights from ATX1 models (x-axis) correlates with those from ATX2 models (y-axis), indicating that similar sets of sequences predict ATX1 and ATX2 localizations. **(e,f)** Clustering and annotation of predictive 6-mers. Each circle represents the top sixty 6-mers with negative (**e**) or positive (**f**) averaged SVM weights in the ATX1 models. Numbers within the circle are ranks of weights’ absolute values (the more ‘predictive’ a 6-mer is, the smaller the number labels it). Pairs of related 6-mers are connected with lines; the thickest lines connect reverse complement pairs, the thinner lines connect neighboring 6-mers with 1 base offset, the thinnest lines connect reverse complement pairs with 1 base offset. Each 6-mers was searched for matching motifs (see Methods), then 6-mer circles were colored corresponding to the category of its top matched motif that meets q-value < 0.1 criteria. 6-mers corresponding to ARGCCCAWT, telobox, GAGA, and TATA-stretch are manually highlighted with border circles. **(g)** Positional distributions of the highly weighted motifs in the TSS region. **(h)** ATX1-bound TSS significantly overlaps with sppRNA-harboring TSS detected in the *hen2-2* background. The significance of the overlap was tested using a hypergeometric test.

**Fig. 5.**
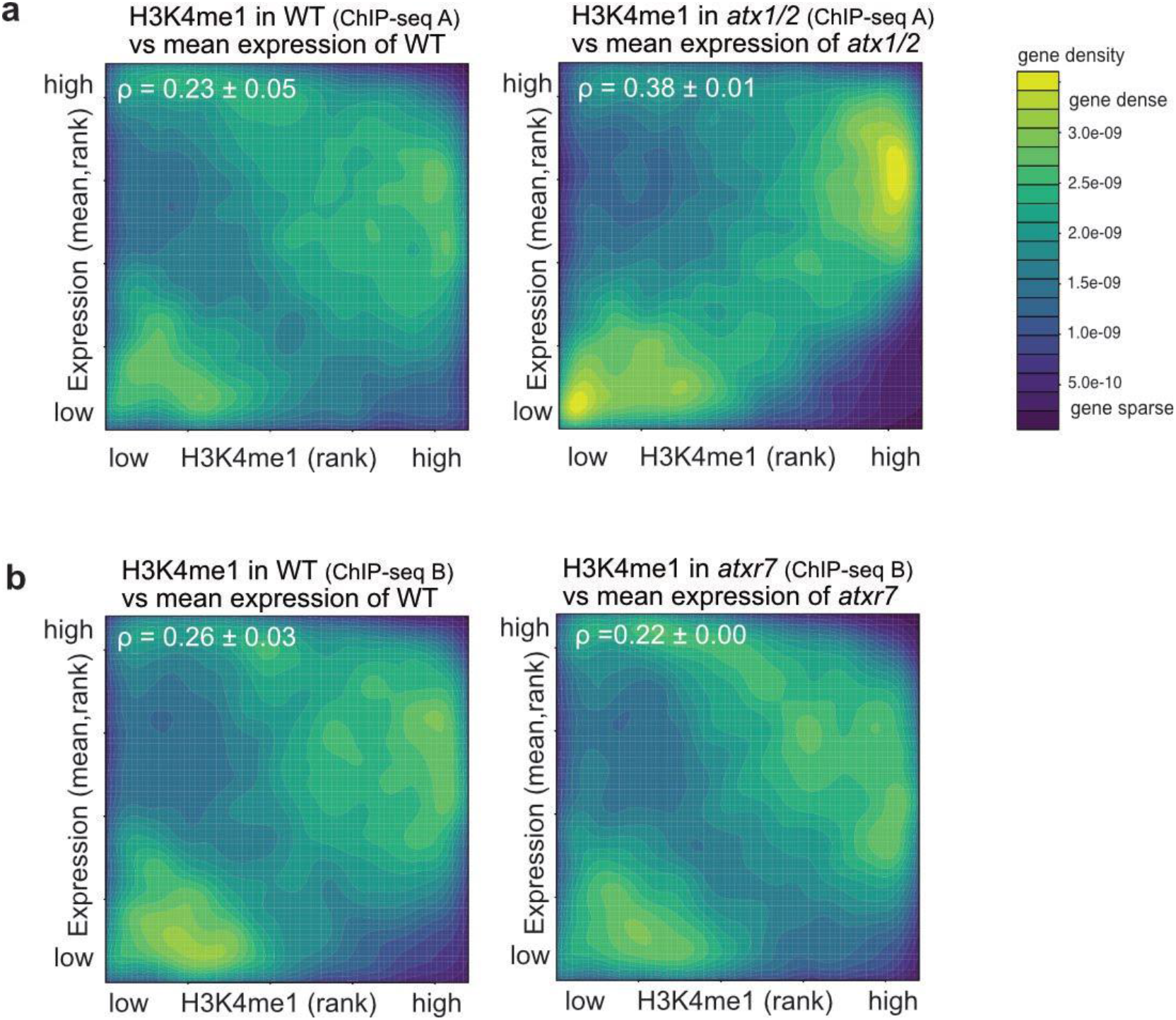
A positive correlation between transcription and H3K4me1 is mediated by ATXR7 and is disturbed by ATX1/2. **(a,b)** Protein-coding genes are ranked in order of H3K4me1 (x-axis, RPKM) and expression levels (y- axis, mRNA-seq FPKM). The densities of genes are visualized as heat maps. ρ is the correlation estimate of Spearman’s test. Three replicates of mRNA-seq resulted in three Spearman’s tests for each genotype, and the average and standard deviation of ρ are represented. The average of the three mRNA-seq results was used for the y-axis. ChIP-seq data used for **(a)** *atx1/2* -WT comparison is the data sets for double mutants shown in Supplementary Fig. 1c (ChIP-seq A), and for **(b)** *atxr7*-WT comparison is the datasets shown in Supplementary Fig. 1a (ChIP-seq B).

The most ‘important’ feature for distinguishing ATXR7-bound genes was the level of RNA Polymerase 2 (RNAP2) around the TTS (Fig. 3a). ATXR7-bound genes showed higher levels of total RNAP2 and RNAP2 phosphorylation at Ser2 and Ser5 sites in the carboxy-terminal domain (CTD) (Fig. 3g, Supplementary Fig. 4), indicating the strong colocalization of ATXR7 and RNAP2, especially when phosphorylated at Ser2 and Ser5. These results suggest that ATXR7, similar to yeast Set1, may be recruited to chromatin in a transcription-coupled manner.

In contrast, the most ‘important’ feature for the prediction of ATX1- and ATX2-bound genes was H3K36me3 (Fig. 3b, c, Supplementary Fig. 4), which gives rise to three nonexclusive hypotheses: the distributions of ATX1/2 and H3K36me3 are driven by a shared factor; the presence of H3K36me3 drives ATX1/2 localization; or ATX1/2 drives H3K36me3 modifications. Consistent with the last hypothesis, *atx1/2/r7* induced the concomitant loss of H3K4me1 and H3K36me3 (Supplementary Fig. 5a), while H3K4me1 was unaffected by the *ash1 homolog 2* (*ashh2*) mutation, which causes a drastic reduction in H3K36me3 levels ^2^^8, 29^(Supplementary Fig. 5b, c). According to the random forest algorithm, the H3K36me3 reduction in *atx1/2/r7* was best explained by the combined H3K4me1 methylation activities of ATX1,2 and ATXR7 (Supplementary Fig. 5d), arguing against H3K36me being directly methylated by some or one of the ATX1,2 or ATXR7. These results suggest that H3K36me3 is regulated downstream of ATX1/2/R7-marked H3K4me1. A mutation in a putative H3K36 methylase, *ASHH3,* affects H3K36me3 at the same target as in ATX1/2/R7, consistent with the view that ASHH3 functions downstream of ATX1/2/R7-marked H3K4me1 to mediate H3K36me3 (Supplementary Fig. 5c, e). In addition to H3K36me3, H4K16ac and H2Bub in TSS regions were indicated to be ‘important’ (Fig. 3b, c). ATX1- and ATX2-bound TSSs were rich in both modifications (Fig. 3h, i, Supplementary Fig. 4). On the other hand, the relative importance values of total or phosphorylated RNAP2 were markedly lower for ATX1/2 than for ATXR7 (Fig. 3a-c). These results suggest that ATX1/2 localization may be governed by other chromatin modification(s), rather than by transcription.

The above results show that there are two modes of chromatin targeting. ATXR7 seems to cooperate with the transcriptional machinery, while ATX1 and ATX2 seem to target genes in a manner that is informed by chromatin modifications, regardless of the abundance of the transcriptional machinery. Interestingly, many genes occupied by ATXR7 are also bound by the H3K4me1 demethylase FLD (^14^, Supplementary Fig. 7a, b). In addition, genes showing the loss of H3K4me1 in the *atxr7* mutant tended to show a gain of H3K4me1 in the *fld* mutant (Supplementary Fig. 7c). Thus, ATXR7 and FLD localize in the TTS region and have opposite effects on H3K4me1.

### The DNA sequences of ATX1- and ATX2-bound TSSs have distinct architectures

Next, we asked whether the DNA sequences to which ATX1,2 or ATXR7 bind also have specific features. We converted the DNA sequences of TSS regions into a vector that represents the occurrence of all possible 6 base sequences (6-mer frequency vector) and trained the linear support vector machine (lSVM) algorithm to distinguish ATX(R)-bound and ATX(R)-unbound regions. This process achieved good accuracy for ATX1 and ATX2 (Fig. 4a, b) but not for ATXR7 (Fig. 4c). The lSVM algorithm learns to assign greater weights to predictively important features. Consistent with their overlapping localization (Fig. 2c), the ATX1 and ATX2 models weighted similar sets of 6-mers (Fig. 4d), suggesting that ATX1 and ATX2 have similar mechanisms of sequence-based targeting.

To better understand the underlying mechanism, we sought to annotate the sets of DNA motifs that specify ATX1/2 binding. We selected the 120 predictive 6-mers, including the sixty 6-mers with the highest positive weights (when the TSS region contains these 6-mers, ATX1/2 are more likely to bind) and the sixty 6-mers with the greatest negative weights (opposing binding). The prediction accuracy for these 120 features was not much lower than that of the full model (Fig. 4a, b). Many of the 120 predictive 6-mers in ATX1/2 models were clustered into groups of overlapping 6-mer sequences (Fig. 4e, f, Supplementary Fig. 10a, b), in contrast to the situation for random or nonpredictive 6-mers (Supplementary Fig. 9a, b, Supplementary Fig. 10c), suggesting the presence of > 6 bp motifs that are predictive of ATX1/2 binding.

We annotated these discriminative DNA motifs by searching for matching sequences in the literature and in databases (Fig. 4 e, f, Supplementary Fig. 10a, b, Supplementary Tables 1, 2). In both the ATX1 and ATX2 models, TATA stretch was negatively weighted (Fig. 4e, Supplementary Fig. 10a, purple circles). Consistent with this result, the occurrence of TATA stretch was less frequent in ATX-bound TSSs than in ATX-unbound TSSs (Fig. 4g, Supplementary Fig. 10d). On the other hand, GAGA stretch was positively weighted (Fig. 4f, Supplementary Fig. 10b, gray circles), and ATX-bound TSSs showed broad GAGA peaks (Fig. 4g, Supplementary Fig. 10d). The positively weighted motifs also included ‘ARGCCCAWT’ (in both the ATX1 and ATX2 models) and a telobox fragment (in the ATX1 model), the latter of which coincided with a trihelix transcription factor (TF) binding site (Fig. 4f, g and Supplementary Fig. 10b, d. Supplementary Fig. 11). RGCCCAW is likely bound by TCP family TFs ^30–32^. While the ATX protein itself is not known to show DNA binding specificity, these additional DNA binders, such as the GAGA binding factor BPC6 ^33, 34^ and TCP and trihelix TFs, may physically link ATX to DNA motifs. The combination of GAGA and RGCCCAW is characteristic of TSSs that harbor short noncoding RNAs (short promoter proximal RNAs, sppRNAs) with unknown functions ^35^. Indeed, ATX1/2-bound genes were found to significantly overlap with sppRNA genes (Fig. 4h, Supplementary Fig. 10e), suggesting the existence of mechanical links between sppRNA and ATX1/2 localization.

These sequence-based learning results further support the idea that ATX1 and ATX2 target genes independent of transcription events. The predictive motif sets are further interpreted in the discussion.

### A positive correlation between transcription and H3K4me1 is mediated by ATXR7 and is disturbed by ATX1/2

From the above results we hypothesized that the ATX1 and ATX2 proteins monomethylate H3K4 irrespective of transcription, while ATXR7 does cotranscriptionally. We tested this hypothesis by analyzing the correlation between transcription and H3K4me1. In the wild type, the amount of H3K4me1 covering a gene is moderately correlated with the transcription level (Fig. 5a, b, left). The hypothesis predicts that ATXR7 is responsible for this correlation through H3K4me1 deposition in transcribed genes, while ATX1 and ATX2 disrupt this correlation by depositing H3K4me1 regardless of transcription. In agreement with this prediction, the strength of the correlation estimated by Spearman’s test or visualized in a density heat map showed that the correlation became stronger in *atx1/2* (Fig. 5a) and weaker in *atxr7* (Fig. 5b) relative to the corresponding wild-type data. This alteration in the correlation landscape was due to a change in H3K4me1 rather than transcription because the correlation showed the same trend when the transcription data for the wild type were used rather than those for the mutants (Supplementary Fig. 12).

### Localization and functional mode of other *Arabidopsis* H3K4 methylases

In addition to ATX1, ATX2, and ATXR7, *Arabidopsis* has at least four other putative H3K4 methylases. Hence, we wondered how the localization patterns of the other methylases were determined. A random forest analysis using the reported genome-wide distribution of the ATXR3 protein ^36^ revealed that the levels of total and phosphorylated RNAP2 and transcription presented the highest ‘importance’ and that they were significantly higher in ATXR3-bound genes than in ATXR3-unbound genes (Fig. 6a-d), suggesting a cotranscriptional mode of H3K4me3 deposition by ATXR3. Consistent with this interpretation, the strong correlation between transcript levels and H3K4me3 decreased in the *atxr3* mutant (Fig. 6e, f, Supplementary Fig. 13a-b).

**Fig. 6.**
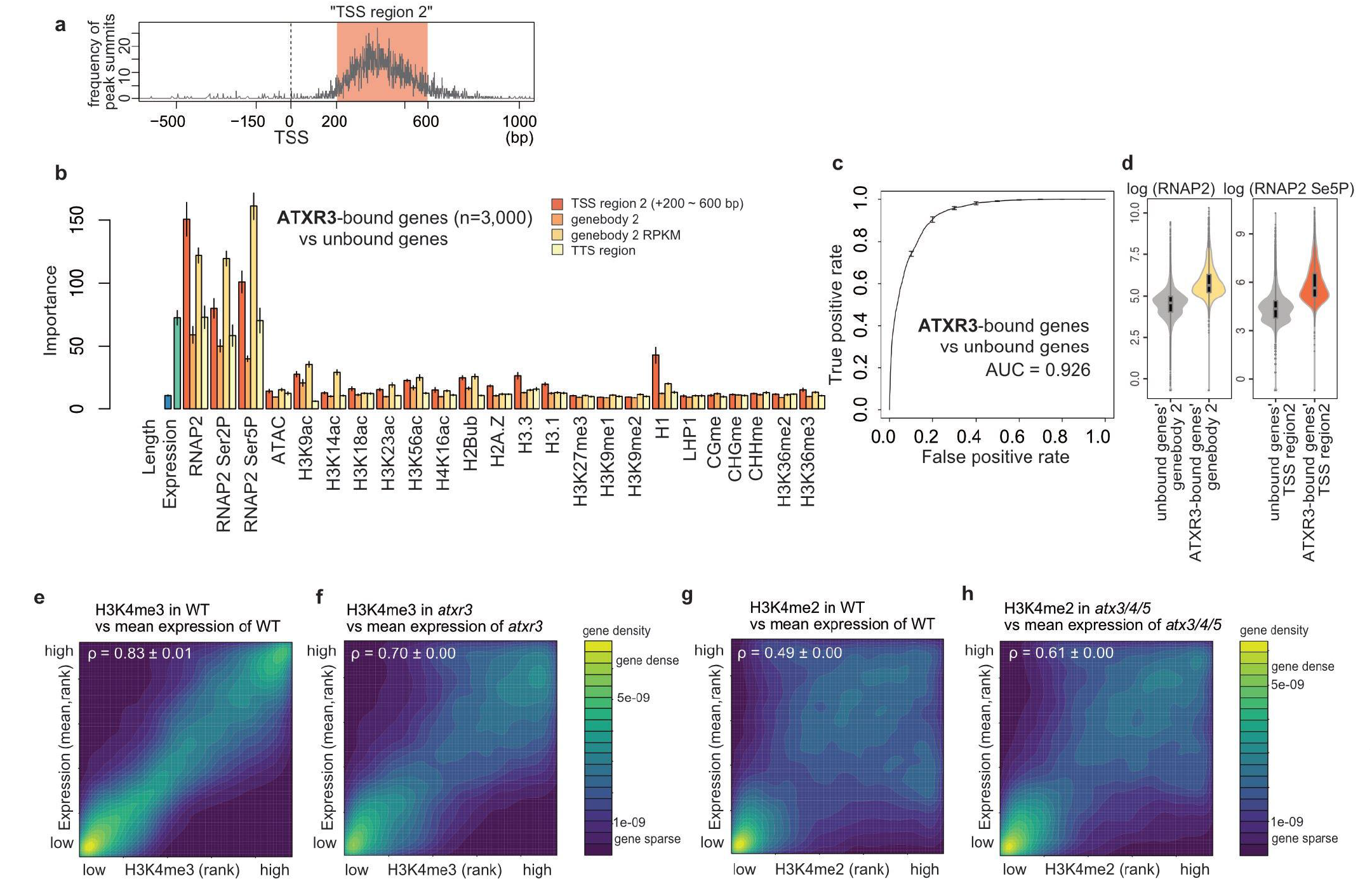
Localization and functional mode of other Arabidopsis H3K4 methylases. **(a)** Position of ATXR3 peaks relative to TSS, visualized as a frequency of ChIP-seq peak summits around TSS. x-axis, distance from TSS; y-axis, number of peak summits. Most of the ATXR3 peaks belong to the region spanning from 200 bp to 600 bp downstream of TSS, which we hereafter refer to as ‘TSS region 2’. **(b)** Chromatin features predictive of ATXR3 localization. Error bars represent the standard deviation of the 5 repeats of training. **(c)** ROC plot showing the prediction accuracy of the random forest models. AUC indicates the area under the ROC curve. ROC and AUC are calculated with data on Chr 5, which are held out from the training as test data. Average and standard deviation of the 5 repeats of training are plotted. **(d)** Violin plots showing the abundance of the two most predictively ‘important’ features. **(e-h)** Protein-coding genes are ranked in order of H3K4me3 **(e, f)** or H3K4me2 **(g, h)** (x-axis, RPM) and expression levels (y-axis, mRNA-seq FPKM). The densities of genes are visualized as heat maps. ρ is the correlation estimate of Spearman’s test. Three replicates of mRNA-seq resulted in three Spearman’s tests for each genotype, and the average and standard deviation of ρ are represented. The average of the three mRNA-seq results was used for the y-axis. ChIP-seq data used is the data sets shown in Fig.1d (ChIP-seq C).

The triple mutation of three other methylases, ATX3/4/5, resulted in H3K4me2 loss (loss was also observed for H3K4me3 in the *atx3-1/4/5* mutant) (Fig. 1d, Supplementary Fig. 2 and ^21^). The genome-wide distributions of the ATX3/4/5 proteins have not been reported. However, the correlation between H3K4me2 and transcription became more prominent in the absence of ATX3/4/5 (as observed for H3K4me3 in the *atx3-1/4/5* mutant) (Fig. 6g, h, Supplementary Fig. 13a, c-i), suggesting that these enzymes introduce H3K4me modifications in a transcription-independent manner, which could be instructed by other chromatin modifications or DNA sequences, as discussed for ATX1/2.

### Localization and functional mode of mammalian H3K4 methylases

We wondered whether these two functional modes of H3K4 methylases also occur in organisms other than plants. Fig. 7a summarizes the lineage of H3K4 methylases in eukaryotes. H3K4 methylases are classified into Set1-type and Trx/Trr-type methylases. ATXR7 and, probably, ATXR3 are Set1-type methylases, while ATX1 to ATX5 belong to the Trx and/or Trr-type methylases (Fig. 7a, ^18, 37^). Fungi have lost Trx/Trr-type genes during evolution and exhibit only one Set1-type gene ^16^. We analyzed mammalian SET1A, MLL2, and MLL3/4, for which genome-wide localization data in mouse ESCs are available. The reanalysis of the ChIP-seq datasets revealed that SET1A ^38^ and MLL2 ^39^ localize around TSSs (typically in the range of -150∼300 bp) and, to a lesser extent, around enhancers (Supplementary Fig. 14a, b). On the other hand, MLL3/4 ^40, 41^ localize to enhancers (typically in the range of -900 ∼ 900 bp from the center of the enhancer) (Supplementary Fig. 14a, b).

**Fig.7.**
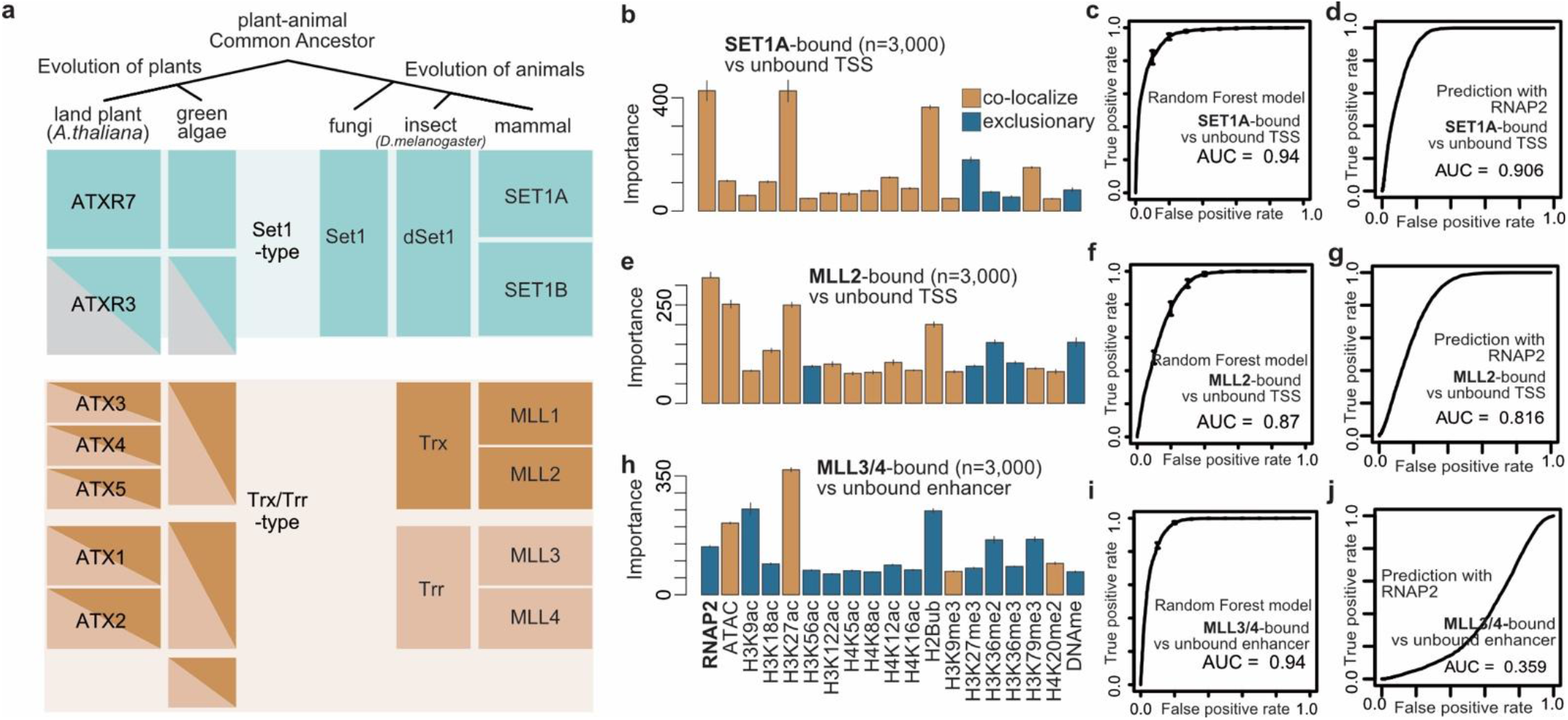
Localization and functional mode of mammalian H3K4 methylases. **(a)** Evolution of H3K4 methylases in eukaryotes based on ^18, 37^. **(b,e,h)** Chromatin features predictive of the localizations of SET1A **(b)** and MLL2 **(e)** at the TSS regions, and MLL3/4 **(h)** at the enhancer regions. Bars are colored brown if the mean abundance of the feature is bound region > unbound region (i.e., colocalize), and colored blue otherwise (i.e., exclusive). Error bars are the standard deviation of the 5 repeats of training. **(c,f,i)** ROC plot showing the prediction accuracy of the random forest models corresponding to **(b,e,h)**. **(d,g,j)** ROC plot showing the prediction accuracy using levels of RNAP2 as the sole predictor. All ROC and AUC are calculated with test data (25% of the original data). Error bars represent the standard deviation of the 5 repeats of training. All the genomic data used here are curated from preks (Supplementary Table 3).vious wor

According to the random forest model, the best predictor of both SET1A-bound TSSs and enhancers was the level of RNAP2 (Fig. 7b, c, Supplementary Fig. 14c, d). SET1A strongly colocalizes with RNAP2 (Fig. 7b, d). MLL2-bound TSSs were also best predicted by RNAP2 (Fig. 7e, f, Supplementary Fig. 14e, f). However, the prediction accuracy for the localization of MLL2 according to the RNAP2 level was lower than that for SET1A (Fig. 7d, g), and RNAP2 was not an outstanding predictor (Fig. 7e), suggesting that the colocalization of RNAP2-MLL2 was weaker than that of RNAP2-SET1A. The best predictor of MLL3/4-bound regions was not RNAP2 but H3K27ac, at both enhancers and TSSs (Fig. 7h, i, Supplementary Fig. 14g, h). At enhancers, which are the major localization sites of MLL3/4 (^40, 41^ Supplementary Fig. 14a, b), the observed localization was exclusive to sites of RNAP2 occurrence (Fig. 7h, j). These results suggest that SET1A is a cotranscriptional H3K4 methylase, as is the case for its *Arabidopsis* homolog ATXR7 (and perhaps ATXR3), while MLL3/4 target chromatin irrespective of transcription, similar to ATX1/2. MLL2 appears to be an intermediate. Therefore, it is plausible that Set1-type H3K4 methylases tend to function in a cotranscriptional manner across eukaryotes, while Trx/Trr-type H3K4 methylases are informed by other chromatin and DNA sequence features (Fig. 8).

**Fig. 8.**
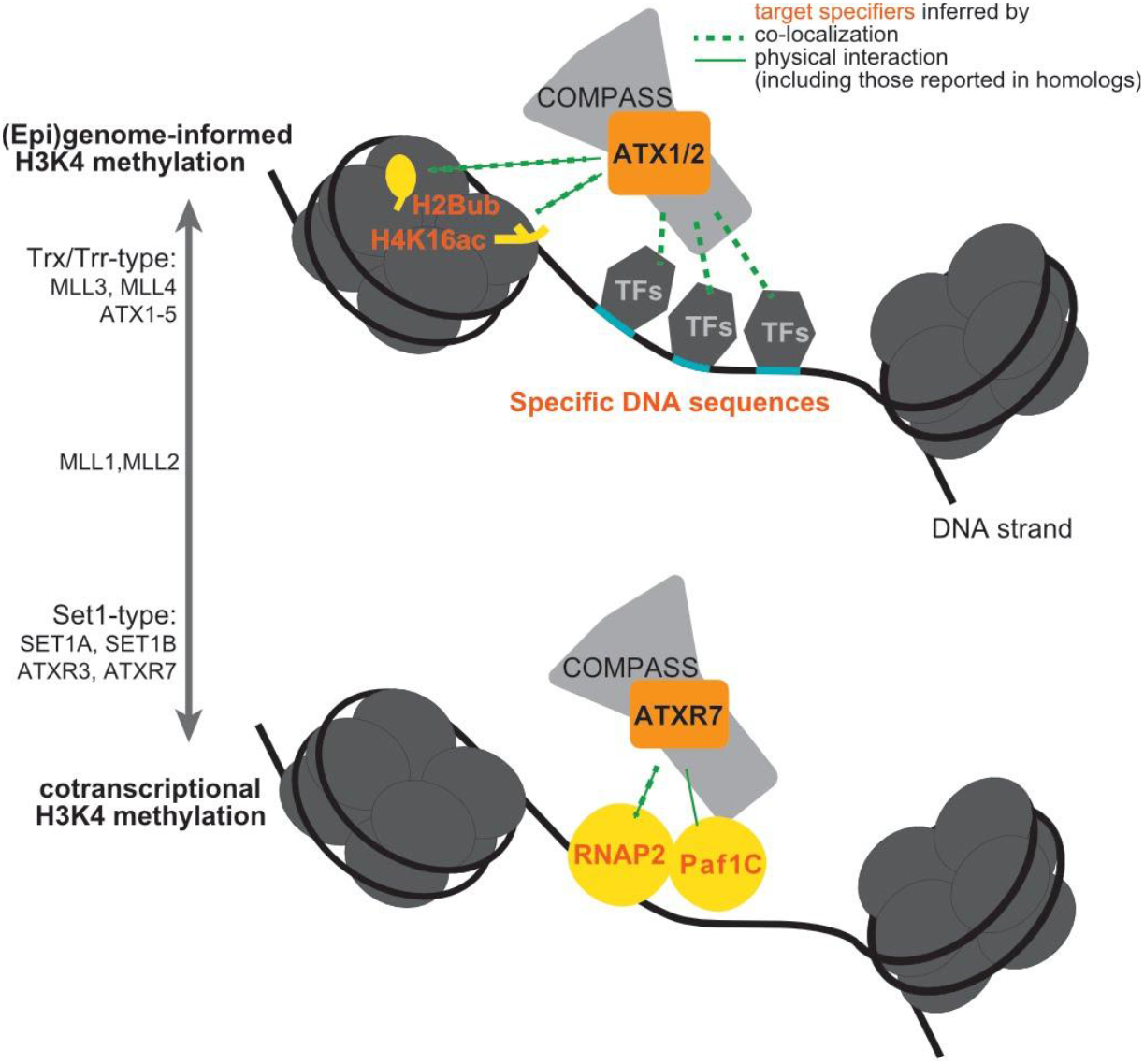
(Epi)genome-informed and cotranscriptional modes of H3K4 methylation.

## Discussion

### Tools for studying the control and function of H3K4me1/2/3

Here, we revealed in *Arabidopsis* that the simultaneous loss of the SET-domain histone methylase genes *ATX1*, *ATX2*, and *ATXR7* causes substantial H3K4me1 loss. In addition, consistent with previous studies ^19–21^, our results showed that ATX3, ATX4, and ATX5 redundantly regulate H3K4me2/3 and that ATXR3 regulates H3K4me3. By using a weak allele of ATX3, we were also able to obtain a mutant with specific H3K4me2 loss. Interestingly, these three groups of SET-domain genes are responsible for H3K4me1, H3K4me2/3, and H3K4me3 modifications among a distinct spectrum of genes. The target genes of these three groups are mutually exclusive, and mutations within them affect H3K4me independently. For example, triple *atx1/2/r7* mutation causes the loss of only H3K4me1 and not H3K4me2/3. This observation implies that other ATX(R)s can catalyze H3K4me2/3 modifications on unmethylated H3K4 (H3K4me0) in their target regions. Accordingly, ATXR3 was shown to catalyze the H3K4me3 from H3K4me0 *in vitro* ^19^. Our study clarified the functional division of labor among ATX(R)s genome wide. In contrast to *Arabidopsis*, yeasts have a single H3K4 methylase, Set1, which is responsible for all H3K4me modifications. The *Arabidopsis* mutants showing the specific reduction of each H3K4me state identified here will serve as powerful materials for understanding the specific functions of H3K4me1/2/3.

Previous works have reported that *atx1* shows H3K4me3 loss ^22–24, 42, 43^ and that *atx2* shows H3K4me2 loss ^23^, on the basis of ChIP-qPCR analyses of selected loci, while our data show that the redundant roles of these enzymes have the largest effect on the global H3K4me1 level. The catalytic domains of ATX1/2/R7 contain a bulky tyrosine at the tyrosine/phenylalanine switch (Supplementary Fig. 15), which is proposed to act as an obstacle to higher-order methylation ^44^, consistent with our conclusion that ATX1/2/R7 primarily regulates H3K4me1. The results of previous studies might reflect indirect effects triggered by altered transcription and/or minor locus-specific effects. Our conclusion is also supported by the results showing that ATX1/2/R7 redundantly repress flowering via *FLC* activation and that ATXR7 and FLD counteract each other in *FLC* regulation, probably by modulating H3K4me1 ^14, 25, 45^.

### Chromatin-targeting mechanisms of ATX(R)s predicted through machine learning analyses

Random forest analyses of the genome-wide localization of ATX1 and ATX2 suggested that H3K36me3, H4K16ac and H2Bub are candidate chromatin features for the recruitment of ATX1/2 (Fig. 3, Supplementary Fig. 4). Among these three candidates, the contributions of H4K16ac and H2Bub to ATX1/2 recruitment agree well with previous reports in other species. MLL3 and MLL4 (mammalian homologs of ATX) directly bind to H4K16ac via plant homeodomains (PHDs) ^46^. ATX1 and ATX2 (but not ATXR7) also have PHD domains (Fig. 1c). The requirement of H2Bub for H3K4me modification has long been known, and various explanatory mechanisms have been proposed. Recently, cryo-EM studies revealed that MLL-type H3K4 methylase-containing complexes (complex proteins associated with Set1, COMPASS) attach more firmly to ubiquitinated nucleosomes ^47^.

Motif mining with the lSVM algorithm revealed three notable characteristics of the ATX1/2-bound TSS sequences that coincided with known features of animal Trx/Trr-bound sequences. First, the ATX1/2-bound regions are GAGA-type promoters rather than TATA-box promoters. In *Arabidopsis*, promoters without TATA boxes (a major core promoter motif) tend to contain GAGA stretch, which are suggested to be equivalent to animal CpG islands ^48^. MLL2 also binds to non-TATA(=CpG)-type promoters ^39^. Second, ATX1/2-bound TSS regions potentially recruit polycomb-group (PcG) proteins, considering that the combination of telobox and GAGA sequences is necessary and sufficient for the recruitment of *Arabidopsis* PcGs ^49^, which are antagonists of trithorax group (TrxG) proteins. In both plants and animals, TrxG and PcG often target the same genomic regions. Third, both ATX1/2-bound and MLL3/4-bound regions produce noncoding RNAs: sppRNAs and enhancer RNAs ^40^. These results suggest that Trx/Trr methylase bind to a common set of features across eukaryotes

### Two modes of methylase recruitment in multicellular eukaryotes

Our analyses suggested that Set1-type methylases typically function in a cotranscriptional manner and that Trx/Trr-type methylases are less informed by transcription. Although this bifurcation has not been explicitly noted previously, it may underlie the observations of a number of previous works. For instance, the concept that H3K4 methylation occurs cotranscriptionally was founded on studies of Set1 in budding yeast ^4–7^. An interactome analysis of RNAP2 recovered Set1-type methylases but not Trx/Trr-type methylases in human cells ^50^, and the colocalization of RNAP2 with Set1-type, but not Trr/Trx-type methylases, is prominent at the microscopic level in the *Drosophila* polytene chromosome ^51^. In contrast, various recruitment factors in addition to RNAP2 have been proposed for Trx/Trr-type methylases, including chromatin modifications, as described above, and long noncoding RNAs ^52^. These genome- and/or epigenome-informed methylases may have provided the foundation for multicellularity via the elaborate control of H3K4me, considering that Trr/Trx-type methylases are present in animals and plants but are absent in yeasts. Regulation by both transcription-coupled and (epi)genome-encoded mechanisms may be further generalized to other chromatin modifications, such as H3K36me ^53^.

In addition to transcription-coupled and (epi)genome-encoded pathways which regulate H3K4 methylases, demethylases also shape the pattern of H3K4me. In *Arabidopsis*, the histone demethylase LDL2 removes H3K4me1 marks from the gene bodies that accumulate H3K9me2 and silences gene expression ^13^. Another histone demethylase, FLD, removes H3K4me1 from sites where convergent overlapping transcription takes place ^14^. Although the biological function of H3K4me remains unclarified, these diverse mechanisms converging on H3K4me1 suggest that the roles of H3K4me require the coordination and integration of a wide range of information. H3K4me is proposed to coordinate splicing ^54^, transcriptional stability ^55^, cryptic transcription ^56^, sense-antisense transcriptions ^14, 57^ and to confer transcriptional memory ^58, 59^. These apparent multifaceted functions may reflect the multiple regulation mechanisms of this modification. In future studies, the dissection of H3K4me based on its controlling mechanisms may help elucidate the functions of H3K4me.

## Materials and Methods

### Plant materials

T-DNA insertion mutants^60, 61^ used in this study were previously described. Names of mutant alleles such as *“atx1-1”* or *“atx1-2”* collide between papers, and the names provided below are not consensus, but examples; *atx1-2* (*sdg27*, SALK_149002C), *atx1-3* (SALK_119016C), *atx2-1* (*sdg30*, SALK_074806C), *atx2-2* (SALK_117262), *atx3-2* (*sdg14*, SAIL_582_H12), *atx3-1* (GK-128H01), *atx4* (*sdg16*, SALK_060156), *atx5* (*sdg29*, SAIL_705_H05), *atxr7-1* (*sdg25*, SALK_149692C), *atxr7-2* (SAIL_446_F12), *atxr3* (*sdg2*, SALK_021008), *ashh1* (*sdg26*, SALK_013895), *ashh2* (*sdg8*, SALK_065480), *ashh3* (*sdg7*, SALK_131218C), *ashh4* (*sdg24*, SK22803), and *ashr3* (*sdg4,* SALK_128444). For *atx1, atx2,* and *atxr7* mutants, *atx1-2, atx2-1,* and *atxr7-1* alleles were mainly used. *atx1-3, atx2-2, atxr7-2* were used as a replicate to confirm bulk H3K4 methylation levels of *atx1/2/r7* mutant in Fig. 1e. For tagged ATX1, ATX2, and ATXR7 lines, the constructs were made with the uniform design; native promoter, which is 3 kb region upstream from initiation codon, followed by 3 x FLAG sequence, coding regions (genomic DNA sequence including introns until just before stop codon) and HA sequence, cloned into pPLV01 vector ^62^. The plasmids were transferred into respective single mutants (ATX1-tag into *atx1-2*, ATX2-tag into *atx2-1,* and ATXR7-tag into *atxr7-1*) via *Agrobacterium tumefaciens* GV3101::pMP90. Plant lines having homozygous transgene were selected in later generations. All the mutants and ‘Wild type (WT)’ are in the Columbia-0 background. For all the experiments, seeds were sown on Murashige and Skoog (MS) plates and kept in dark at 4°C for a few days, then grown for 15 days under long-day conditions (8 h dark and 16 h light) at 22°C. Whole seedlings were used for experiments.

### ChIP-seq/ATAC-seq/BS-seq

ChIP targeting histone modifications, RNAP2, and epitope-tagged proteins were carried out as described previously ^14^. Antibodies are H3 (ab1791; Abcam), H3K4me1 (ab8895; Abcam), H3K4me2 (ab32356; Abcam), H3K4me3 (ab8580; Abcam), H3K36me2 (MABI0332; MBL), H3K36me3 (MABI0333; MBL), H3K27me3 (MABI0323; MBL), RNAP2 total CTD (CMA601; MBL), RNAP2 phospho S2 (CMA602; MBL), RNAP2 phospho S5 (CMA603; MBL), FLAG (F1804; SIGMA). For ATAC, nuclei preparation and tagmentation procedure followed ^63^ without modification. ChIP-seq and ATAC-seq libraries were made with KAPA Hyper Prep Kit (Kapa Biosystems), and dual size-selected using Agencourt AMPure XP (Beckman Coulter) to enrich 200 – 500 bp fragments. The libraries were 50-bp single-end sequenced by HiSeq4000 sequencer (Illumina) in Vincent J. Coates Genomics Sequencing Laboratory at UC Berkeley. ChIP targeting epitope-tagged proteins and their controls were carried out twice as biological replicates from independently grown plants. Whole-genome bisulfite sequencing (BS-seq) was conducted as described before ^64^. The libraries were 150 bp paired-end sequenced by the HiSeqX Ten sequencer (illumina).

### RNA-seq

Total RNA was extracted from seedling with RNeasy Plant Mini Kit (Qiagen), then polyA selected, fragmented, and made into a library with KAPA Stranded mRNA-seq Kit (Kapa Biosystems) following the manufacturer’s protocol. The libraries were 50-bp single-end sequenced as described above. One sample corresponds to one individual seedling, and three samples were sequenced for one genotype.

### Genomes and Annotations

For *Arabidopsis*, TAIR10 reference genome and Araport 11 gene annotations were used. In this paper, ‘gene’ otherwise stated refers to all the transcribed features in Araport11 annotation, including TE genes or non-coding RNAs. ‘protein_coding’ genes refer to the nuclear genes that are annotated as such in Araport11 annotation, excluding the ones that lack primary transcript annotation. For *Mus Musculus*, we used reference genome mm9, which is not the latest, for compatibility reasons. Gene annotations are from NCBI RefSeq. Enhancers are from Enhancer Atlas V2 ^65^. Corresponding to the cell types of the data origin, MLL3/4 data was analyzed with R1 enhancer annotation, while MLL2 and SET1A data were analyzed with V6.5.

### Sources of reanalyzed data

For *Arabidopsis*, the genomic data were reanalyzed from the following works; H3K9me2,DRR023251^13^; H3K9ac,GSM701925 and H3K18ac,GSM701927 ^66^; H3.1,GSM856055 and H3.3,GSM856054 ^67^; H1,GSM2544793 and H2A.Z,GSM2544791 ^68^; H3K14ac GSM2051285 ^69^; Like-Heterochromatin Protein 1, GSM2028108 ^70^; H3K56ac, GSM2027818 ^71^; H3K23ac, GSM1701017 and H4K16ac, GSM1701018 ^72^; and H2B ubiquitination, GSM3092016 ^73^.

For *Mus Musculus*, the genomic data were reanalyzed from the following works; SET1A ^38^, MLL2 and its control ^39^, MLL3/4, its control, RNAP2, ATAC and H3K27ac ^40^, H4K16ac ^74^, H3K36me2 ^75^, H3K9ac, H3K56ac, H3K36me3 and H3K79me3 ^76^, H4K12ac, H3K9me3 and H4K20me2 ^77^, H3K18ac, H3K122ac, H4K5ac, H4K8ac and H3K27me3 ^78^, and DNAme ^79^. The exact names of the analyzed files are provided in Supplementary Table 3. All data is from mESC.

### Mapping and counting coverages

For *Arabidopsis*, ChIP-seq and ATAC-seq data were processed essentially as described in ^14^. Briefly, our sequenced data were mapped to the TAIR10 reference genome using bowtie ^80^ -v 2 -m 1 option. Mapped reads were extended to 250 bp to represent sequenced fragments, then counted for coverage over the region of interest specified as bed files, using SAMTools ^81^ and BEDTools ^82^. The coverage was RPM/RPKM normalized with custom scripts. Metaplot profiles and heat maps were generated with ngs.plot ^83^. mRNA-seq data were also processed as described in ^14^ using STAR ^84^. BS-seq data were processed as described in ^13^ using Bismark ^85^. For the reanalyses of the published data, SRA data were adapter trimmed following the original papers’ procedure, mapped, and counted essentially the same as above.

For *Mus Musculus*, SRR data of ChIP-seq targeting H3K4 methylases were adapter-trimmed and mapped to mm9 reference genome with -v 2 -m 1 option, then counted and normalized as described above. The other genomics data of *Mus Musculus*, namely the data of chromatin modifications used for the random forest, were curated and parsed from bigWig format provided as supplementary files at Gene Expression Omnibus (GEO) (Supplementary Table 3). bigWig files were first converted to bedGraph with bigWigtoBedGraph ^86^ of UCSC tools, then to bed file with the awk command of Unix, and when applicable, genome version was converted into mm9 with liftOver of UCSC tools, then coverage was counted over the regions of interest.

### Clustering of the 6 single mutants

We selected the top 3,000 genes that had the highest reduction of H3K4me1 (RPKM) compared to WT for each mutant. The regions from TSS to TTS of the 3,000 genes were split into 6 bins. For the WT and the six mutants, the number of reads mapped to each of the 6 sub-gene regions were counted, RPM normalized, and the difference between WT and the mutant was calculated. To capture the trend of which ones of the 6 bins lose H3K4me1, 6 bins in each one gene were standardized. Thus, for each mutant, a matrix of 6 regions x 3,000 genes was obtained. These matrices were vectorized, and Euclid distances were calculated, which were used for hierarchical clustering (Fig. 1b). To explain the idea behind this clustering, considering if a hypothetical methylase catalyzing H3K4me1 into H3K4me2 is lost, H3K4me1 level would be elevated in the region that H3K4me2 originally occupied. We reasoned that similarity in affected sub-gene regions reflects the similarity in function.

### Peak calling

For *Arabidopsis*, enrichment peaks of H3K4 methylases were identified by comparing against negative control (ChIP-seq done with FLAG antibody in non-transgenic WT) using MACS2 ^87^ with option -g 1.3e8 -q 0.3 –nomodel. The distances of the summits of the called peaks from the nearest TSS (in the cases of ATX1, ATX2, and ATXR3) or TTS (in the case of ATXR7) were measured and visualized as a histogram (e.g. Fig. 2d). Based on the histogram, we identified the region that ATX1, ATX2 typically occupy to be 150 bp upstream and 300 bp downstream of TSS, which we named ‘TSS region’. Similarly, ‘TTS region’ is defined as 200 bp upstream to 200 bp downstream of TTS, where ATXR7 localizes, and ‘TSS region 2’ starts from 200 bp downstream of TSS to further 400 bp, where ATXR3 localizes.

For *Mus Muscles*, enrichment peaks of H3K4 methylases were called with default options of MACS2. The distances of the summits of the called peaks from the nearest TSS or the midpoint of enhancer region (in the cases of MLL2 and SET1A, V6.5 enhancer, while in the case of MLL3/4, R1 enhancer) were measured and visualized as a histogram. Based on the histogram, we identified the region that SET1A and MLL2 typically occupy to be 150 bp upstream and 300 bp downstream of TSS, coinciding with the ‘TSS region’ of *Arabidopsis*. Similarly, we defined ‘enhancer region’, where MLL3/4 localizes, to be 900 bp on both sides from the center of Enhancer Atlas’s annotation.

### Random forest

For ATX1, ATX2, and ATXR7 models, genes shorter than 500 bp were filtered out. Gene was divided into three regions; TSS region, TTS region (definitions described above), and gene body region. The gene body region is the transcribed region minus TSS and TTS regions. Similarly, for ATXR3 models, the transcribed regions of genes longer than 800 bp were divided into TSS, TSS2, TTS, and gene body2 regions. Gene body2 region is the coding region minus TSS, TSS2, and TTS regions.

Explanatory features are levels of chromatin modifications/chromatin states calculated over these three regions per gene, using raw read files reported by preceding works (see ‘sources of the reanalyzed data’ section) and this study. For gene body and gene body2 regions, both length-normalized and unnormalized values are included. These data combined with expression level (FPKM of mRNA-seq in WT, from this work) and gene length were used as the predictor variables.

Objective variables were set as two classes; ATX(R)-bound and -unbound genes. ATX(R)-bound genes are defined as the top 3,000 genes that have the highest ChIP-seq signal (RPM values of ATX(R)-Tag expressing lines - negative control) in the typical occupying region defined from the above-mentioned peak calling; TSS region for ATX1 and ATX2, TTS region for ATXR7, TSS2 region for ATXR3. ATX(R)-unbound genes are the rest of protein coding genes (n = all protein coding genes longer than 500 or 800 bp - 3,000). We uniformly set the number of bound genes to 3,000, at the potential expense of predictive accuracy, in order to reduce parameter variance between models.

To balance the two classes, 3,000 genes out of the ATX(R)-unbound genes were randomly selected, and the data on chromosome 1 to 4 were used for training random forest (R package ‘RandomForest’), while data on chromosome 5 were held out as test data.

For SET1A, MLL2, and MLL3/4 models, explanatory features are levels of chromatin states calculated over TSS regions or enhancer regions, using previously reported data in mESC (see ‘sources of the reanalyzed data’ section). Objective variables were set in the same manner as ATX(R)s models; that is, Set1/MLL2/MLL3/4-bound regions were defined as the top 3,000 regions with the highest TSS region signals regions or enhancer regions, and unbound regions were down-sampled into 3,000. 25% of the data were held out from the training and used as test data.

Both for *Arabidopsis* and *Mus muscles*, the random sampling and training steps were repeated 5 times, and the average and the standard deviation of the importance derived from the 5 models were plotted. ROC was plotted based on the prediction of the 5 models, using test data.

### Support Vector Machine

Application of SVM to DNA sequence essentially followed previous work on human enhancer classification ^88^. Briefly, DNA sequences on the coding strand of TSS regions were converted to k-mer frequency vectors of 4^k features. The frequency vectors were standardized and given bound or unbound labels in the same way as random forest.

The data on chromosome 5 were held out as test data. Data on the chromosome 1 to 4 were used for 5 cross-fold validation for training to select the optimal parameters. For each trial of cross-fold validation, data of the unbound class were randomly sampled independently to balance the two classes. The averaged weights of features (k-mer sequences), which were calculated based on the 5 independently trained models using the best parameters, were used for further motif annotations. ROC was plotted based on the prediction of the 5 models with test data. We trained 5,6,7-mer lSVM or 6-mer kernel-SVM to find that the accuracy of 6-mer lSVM was about the same or better than the rest (Supplementary Fig. 16).

### Motif annotation

The decision function of linear SVM is the sum of weighted features (in this case, standardized occurrence of each k-mers), which allows us to interpret the highly weighted motifs to be predictively important ones. Using the weights averaged from the 5 best models, sixty 6-mers that had the largest positive weights, largest negative weights, and smallest absolute weights, and sixty random 6-mers were selected.

To make clusters of related motifs within each group of sixty 6-mers, similarity scores were calculated between all pairs of the 6-mers. Similarity scores are defined as ; reverse complement (= 1) > 1 nt offset > 1 nt offset on the opposite strand > the others (= 0). The resulting similarity score matrices were visualized with the qgraph ^89^ package in R.

Next, to annotate the 6-mers while taking into account the context the 6-mer is located, we calculated the ATGC frequency in the flanking three bases on both sides of each 6-mer. The flanking ATGC frequency of the positively weighted sixty 6-mers was calculated with the sequence of ATX-bound TSS region, while negatively weighted and random/near-zero weighted 6-mers were calculated with the sequence of ATX-unbound and random TSS region, respectively. The resulting probability matrices representing 3 nt + 6 nt + 3 nt were converted into meme format and searched for matching motifs using TOMTOM against all the *Arabidopsis* databases included in the MEME Suite’s motif_databases.12.19 ^90^. For each motif, TOMTOM hits were summarized in Supplementary Table 2, and top-hit satisfying q.value< 0.1 is indicated by colors representing TF families (e.g. Figure. 4e, f). The distribution of the motifs of interest, i.e., GAGA-stretch, TATA-box, telobox, and RGCCCAW, was visualized by the presence of the 6-mers comprising the motifs. Namely, {AAGAGA, GAGAGA, AAGAGG, GAGAGG, AGAGAA, GGAGAA, AGAGAG, GGAGAG} represented GAGA-stretch, {TATAAA, TATATA, ATATAA, ATAAAT, TAAATA, ATATAT, TTATAA, TTATAT} represented TATA, {AAACCC, AACCCT, ACCCAA, ACCCTA} represented telobox and {AAGCCC, AGGCCC, GGCCCA, AGCCCA, GCCCAA, GCCCAT, GGGCTT, GGGCCT, TGGGCC, TGGGCT, TTGGGC, ATGGGC, AATGGG, CCCATT} represented ARGCCCAWT.

### Correlation between H3K4me and transcription

Spearman correlation between mRNA-seq and ChIP-seq data of H3K4me was calculated using all protein-coding genes. mRNA-seq data was FPKM normalized, ChIP-seq data targeting H3K4me1 was RPKM, H3K4me2/H3K4me3 was RPM normalized. For color-filled contour plots, genes for which no transcription was detected were excluded.

### Data availability

The high-throughput sequencing data generated in this study is available in the NCBI database for reviewers during the review process, and for public upon publication.

### Code availability

Codes will be available on GitHub upon publication.

### Author Contributions

S.O., T.K. and S.I. conceived the study. S.O., M.T., K.T., and S.I. performed the experiments. S.O conducted the analysis. S.O. drafted and S.O., T.K. and S.I. edited the manuscript.

## Acknowledgements

We thank all Kakutani laboratory members for helpful discussion and technical assistance. This work used the Vincent J. Coates Genomics Sequencing Laboratory at UC Berkeley, supported by NIH S10 OD018174 Instrumentation Grant. The computations were partially performed on the NIG supercomputer at NIG, Japan. We thank NASC/ABRC for distributing the seeds. The *ashr3* mutant was kindly gifted by Taku Sasaki. This work was supported by grants from Japan Science and Technology Agency (JST) PRESTO (no. JPMJPR17Q1) to S.I., JST CREST (no. JPMJCR15O1) to T.K., and Japan Society for the Promotion of Science (JSPS) (nos. 26221105, 15H05963 and 19H00995 to T.K., 20H05913 to S.I. and 19J21882 to S.O.). S.O. is supported by the JSPS Research Fellowship for Young Scientists.

---

**Supplementary Fig. 1.**
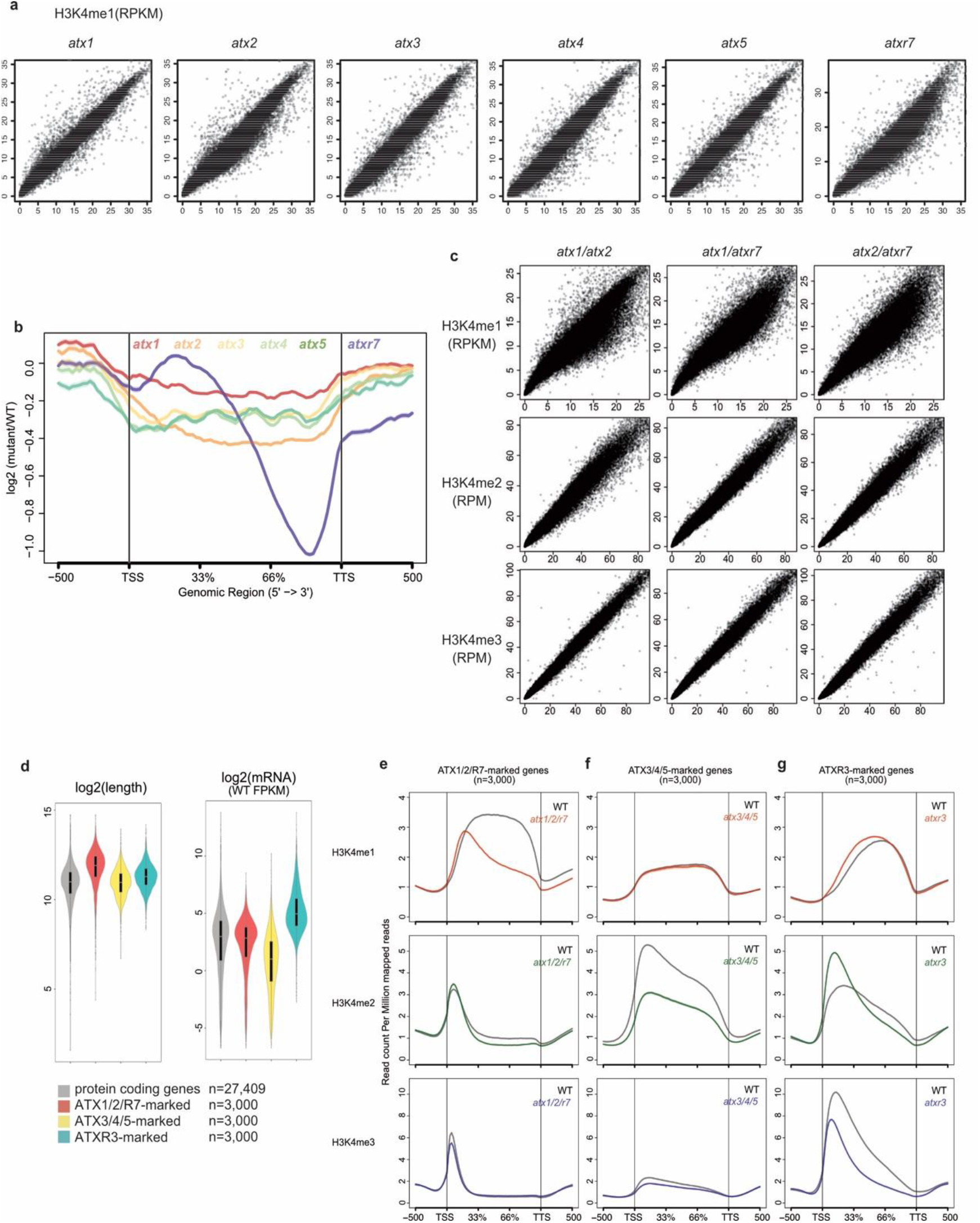
Impacts of *atx(r)* mutations on H3K4me. **(a)** ChIP-seq of H3K4me1 in 6 mutants (y-axis) did not show large change compared to WT (x-axis). Each dot represents one gene and axes are trimmed to 0.98 quantiles. **(b)** Metaplot illustrates the averaged pattern of H3K4me1 changes of the 6 mutants compared to WT. **(c)** ChIP-seq for H3K4me1, H3K4me2, H3K4me3 in *atx1/2, atx1/atxr7 and atx2/atxr7* double mutants (y-axis) compared to WT (x-axis). For all three mutants, H3K4me1 shows the most prominent reduction. **(d)** ATX1/2/R7-, ATX3/4/5-, and ATXR3-marked genes have distinct characteristics in gene length and expression level. **(e)-(g)** Metaplots illustrating the averaged distribution of H3K4me1,2,3 over ATX1/2/R7-marked **(e)**, ATX3/4/5-marked **(f)**, and ATXR3-marked genes **(g)**.

**Supplementary Fig. 2.**
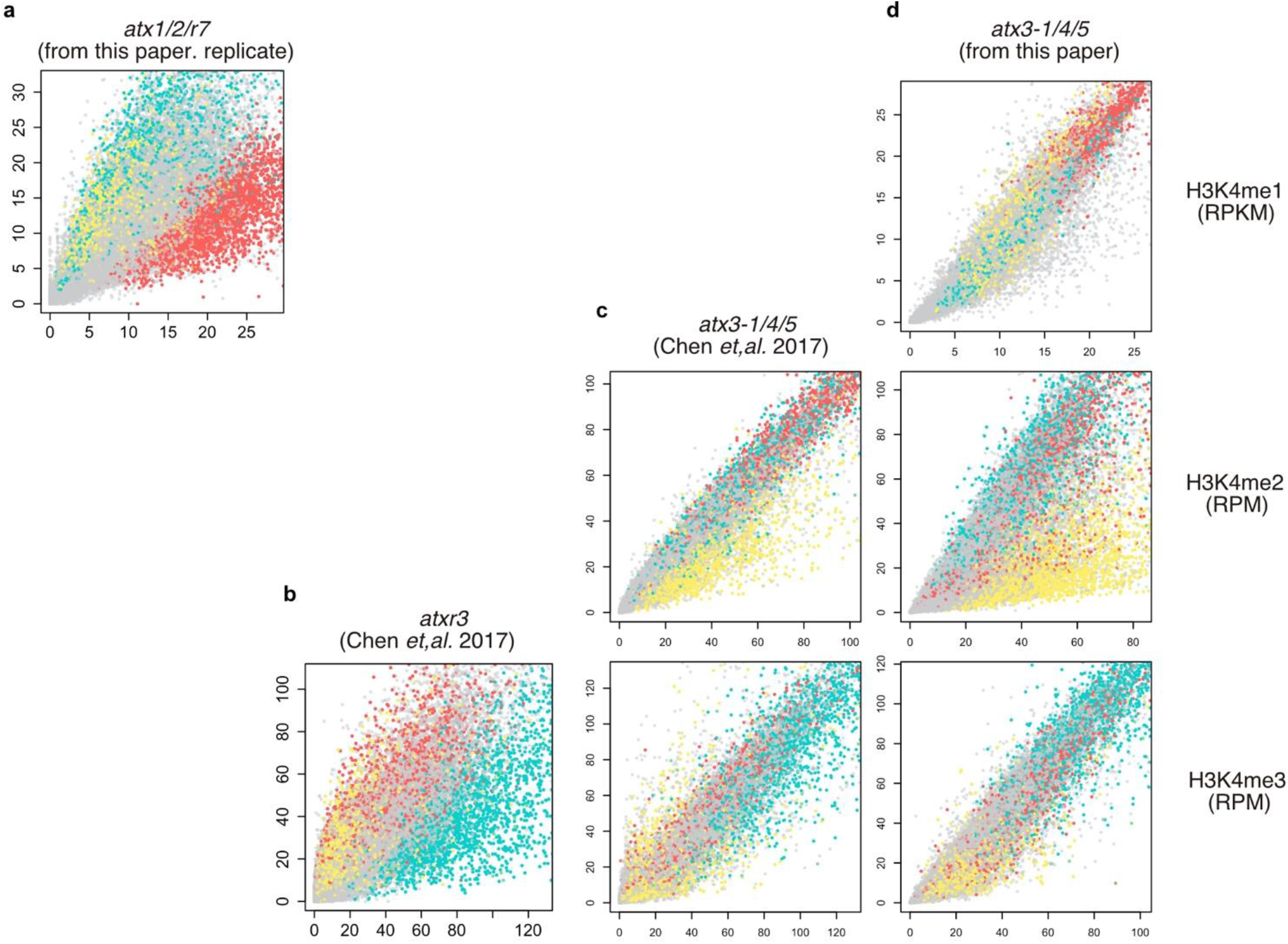
Effects on H3K4me in *atx3/4/5* and *atxr3* mutants agree with previous reports. Mutant (y-axis) ∼ WT (x-axis) comparison of biological replicate of *atx1/2/r7* (**a**), and previously reported ChIP-seq datasets in *atxr3* (**b**), *atx3-1/4/5* (**c**) mutants, and our datasets on *atx3-1/4/5* (**d**). Red, yellow, and blue dots represent ATX1/2/R7-, ATX3/4/5-, and ATXR3-marked genes (see Fig. 1d). Our data show consistent trends with datasets from^21^; in both datasets, *atxr3* mutants lose H3K4me3 in blue-colored genes while *atx3(-1)/4/5* mutants lose H3K4me2 in yellow-colored genes. Axes are trimmed to 0.98 quantiles.

**Supplementary Fig. 3.**
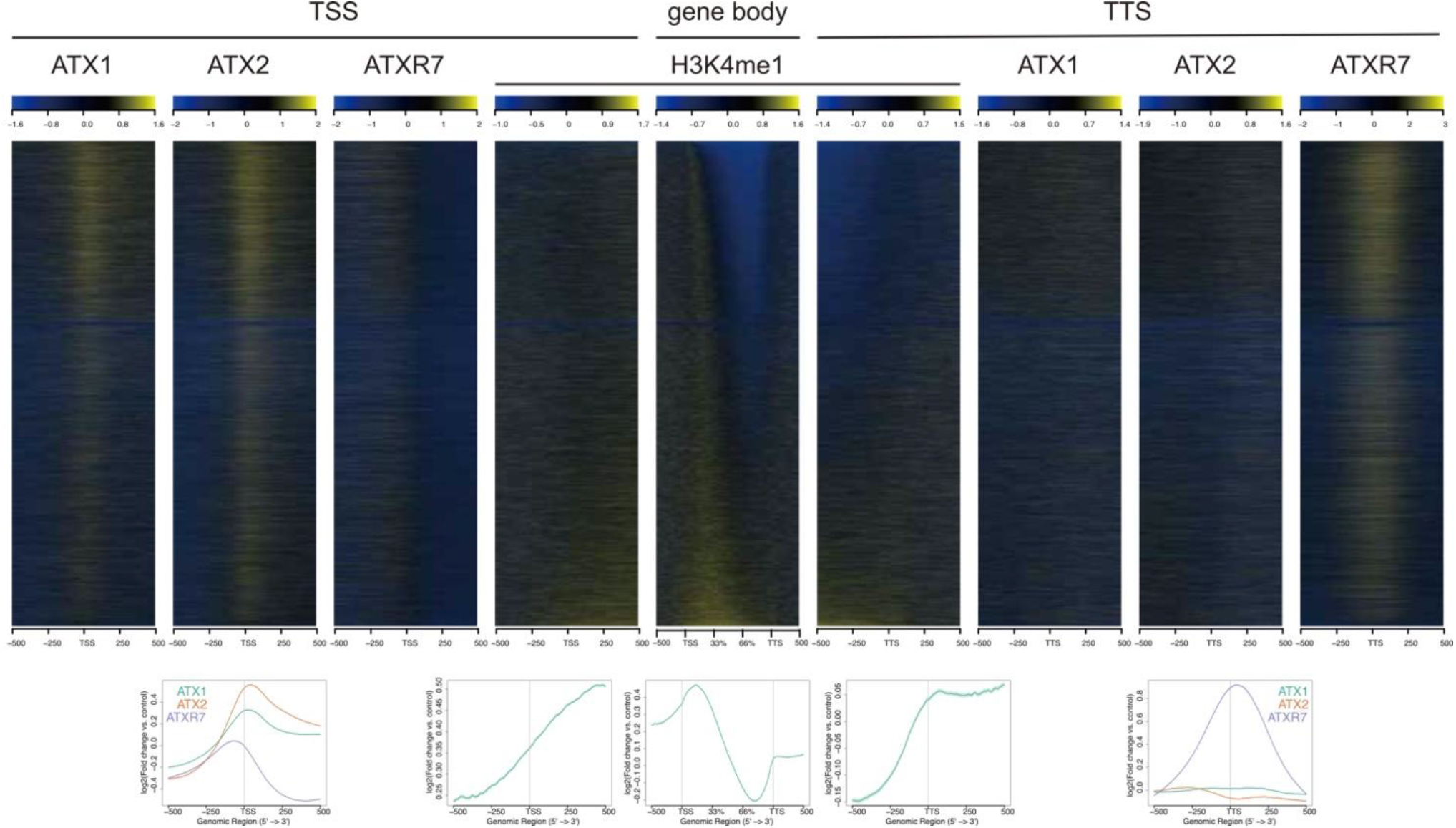
ATX1, ATX2, and ATXR7 occupied genes tend to lose H3K4me1 in *atx1/2/r7*. Metaplots and heat maps showing H3K4me1 change in *atx1/2/r7* and ATX(R)s distribution. All heat maps are sorted according to the H3K4me1 change in *atx1/2/r7* (the heat map in the middle is the same data as Fig.1c, shown for reference) so that genes that lost H3K4me1 the most come to the top.

**Supplementary Fig. 4.**
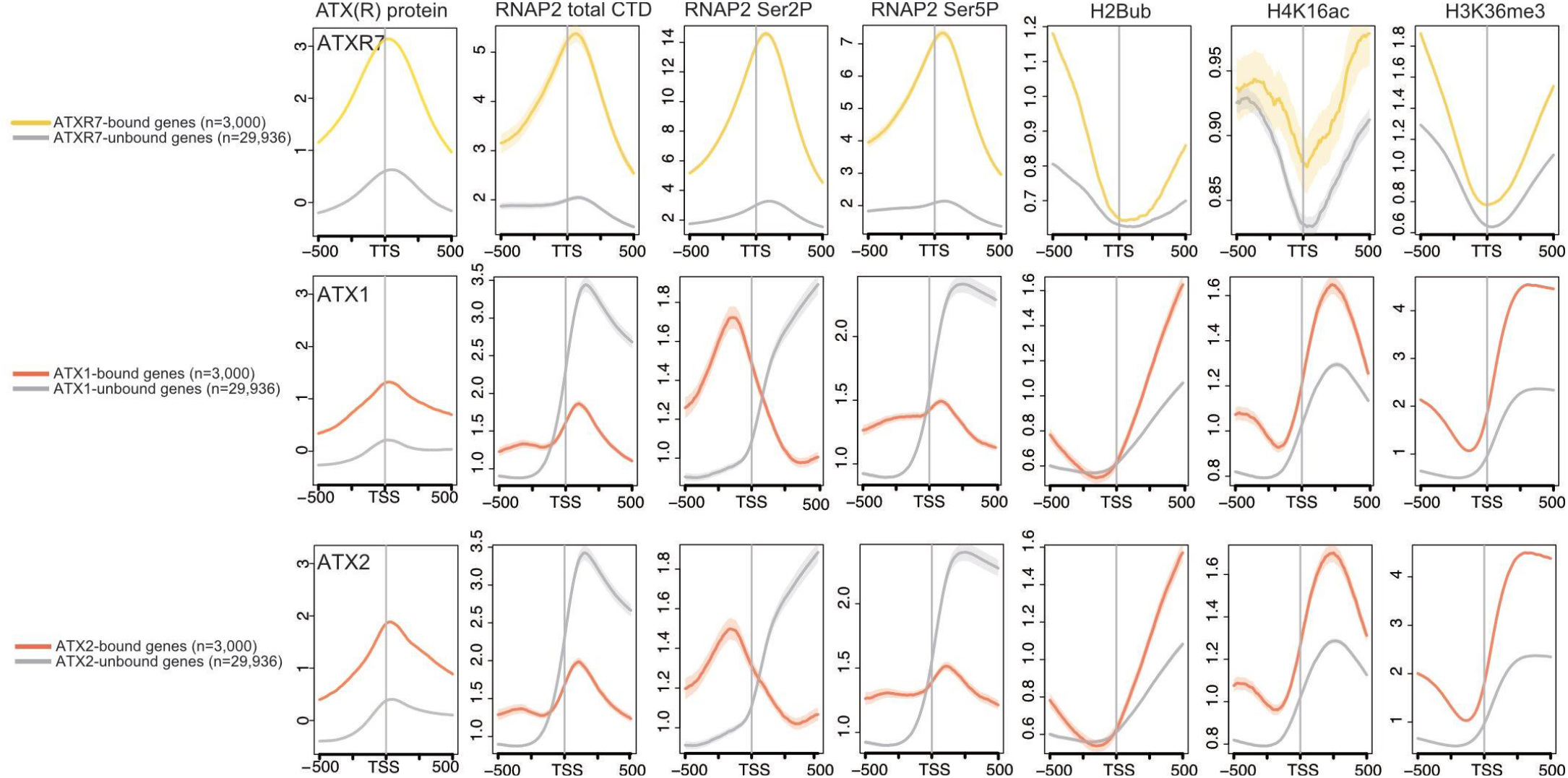
Distribution profiles of predictively ‘important’ chromatin modifications for ATX1, ATX2, or ATXR7 localization. Averaged distribution profiles of ATX(R)s proteins and features that had high importance in the random forest models to predict ATXR7 or ATX1,2-bound genes (see Fig. 3). Profiles around TTS are shown for ATXR7, and profiles around TSS are shown for ATX1 and ATX2.

**Supplementary Fig. 5.**
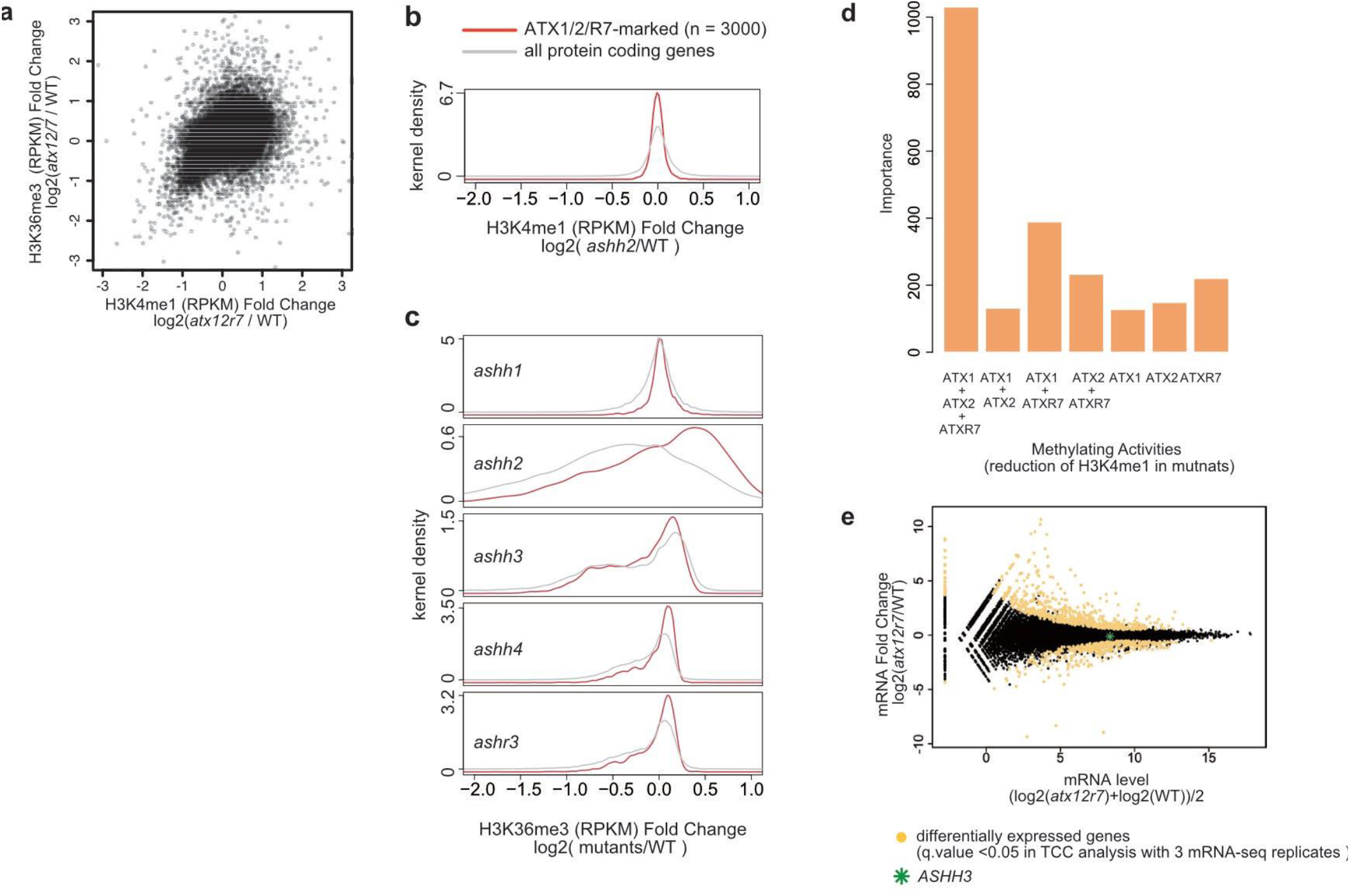
H3K4me1 may promote H3K36me3 via ASHH3. **(a)** H3K36me3 reduction in *atx1/2/r7* (y-axis) correlates with H3K4me1 reduction (x-axis). **(b)** H3K4me1 is not largely affected in the *ashh2* mutant. **(c)** In search for a methylase that catalyzes H3K36me3 in response to ATX(R)s-catalyzed H3K4me1, we analyzed H3K36me3 change by ChIP-seq in mutants of ASHH2, known major H3K36me3 methylase ^28, 29^ and ASHH1, ASHH3, ASHH4, and ASHR3, whose catalytic domain SET are similar with ASHH2 ^17, 18^. Density histograms of H3K36me3 reduction in each mutant show *ashh3* lose H3K36me3 at ATX1/2/R7-marked locus, suggesting that ASHH3 is at least partly responsible for H3K36me3 reduction in *atx1/2/r7*. **(d)** Random forest was trained to predict H3K36me3-reduced region in *atx1/2/r7* based on H3K4me1 reduction in gene body regions of the *atx1*, *atx2*, *atxr7*, *atx1/2*, *atx1/r7*, *atx2/r7,* and *atx1/2/r7* mutants. **(e)** mRNA expression of *ASHH3* was not altered in *atx1/2/r7*, precluding the possibility that H3K36me3 change in *atx1/2/r7* was due to reduced expression of *ASHH3*.

**Supplementary Fig. 6.**
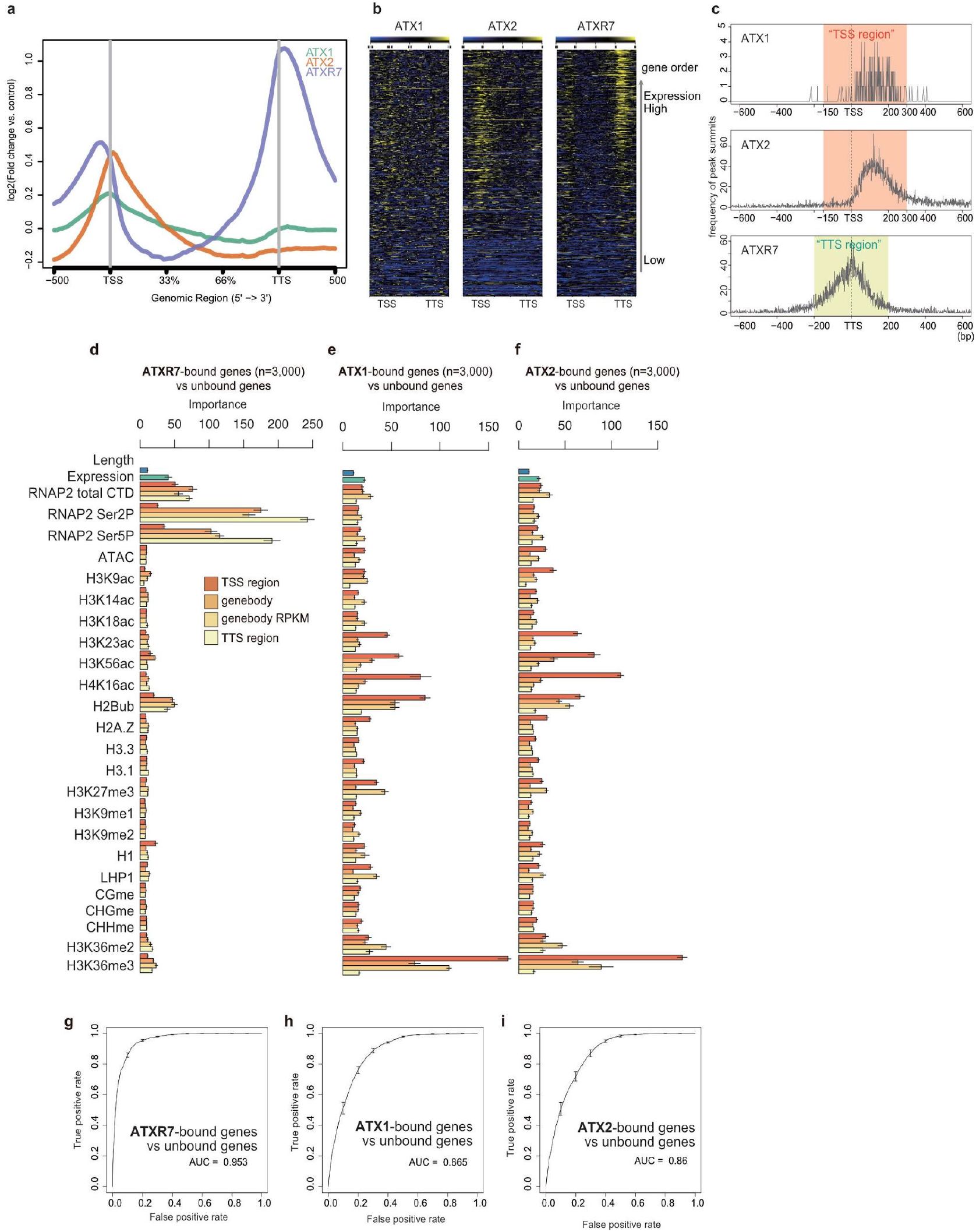
Random forest analyses using the biological replicates of ATX(R)s ChIP-seq. The ChIP-seq data sets of the biological replicates, which were independently grown, were processed in the same manner as in Fig. 2 and Fig. 3. **(a)** Metaplot and **(b)** heat maps illustrating ATX1, ATX2, and ATXR7 distributions in the gene body region corrected with non-transgenic control. The heat maps were sorted so that highly transcribed genes (measured as mRNA-seq in WT) come to the top. **(c)** Position of ATX1 and ATX2 ChIP-seq peaks relative to TSS and ATXR7 peaks relative to TTS (x-axis), visualized as a frequency of peak summits (y-axis). Consistent with the other replicates (Fig. 2), most of ATX1 and ATX2 peaks belong to the ‘TSS region’, while most of ATXR7 peaks belong to the ‘TTS region’. **(d-f) ‘**Importance’ of features and **(g-i)** ROC plot of random forest model trained to predict ATX1/2/R7-bound genes. Average and standard deviation of the 5 repeats of training are plotted.

**Supplementary Fig. 7.**
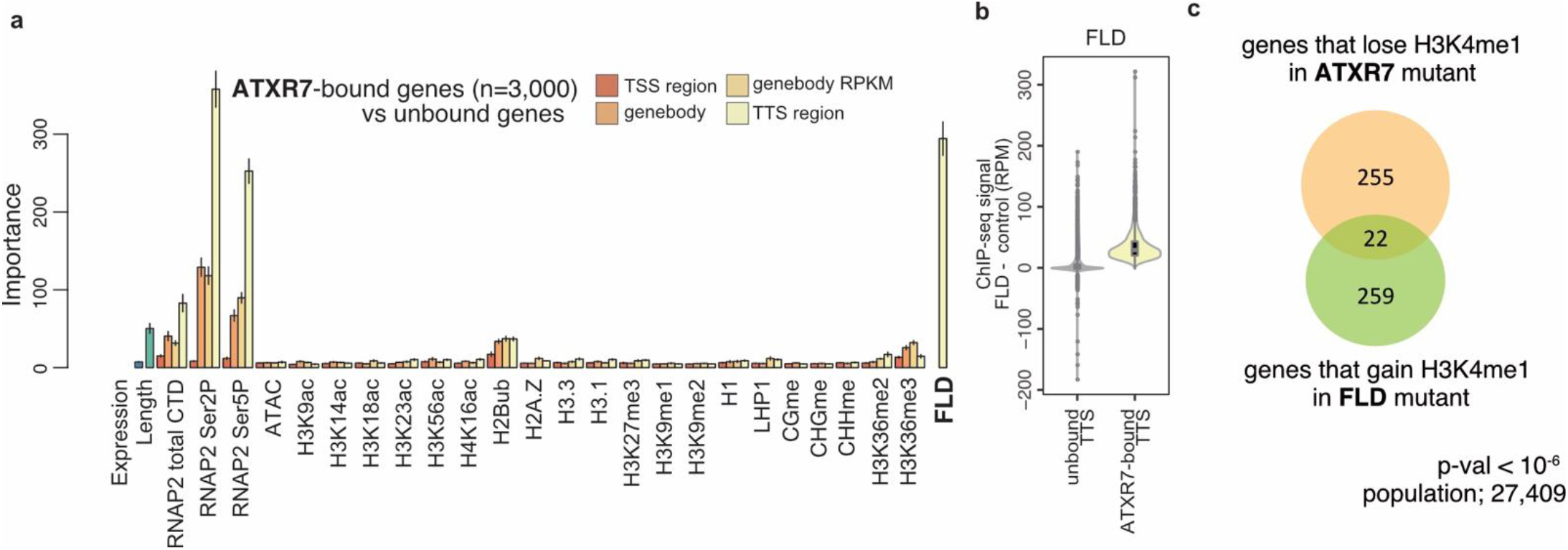
ATXR7 and FLD share target genes. ATXR7 and FLD colocalize. When the Random Forest model is trained to predict ATXR7-bound and -unbound genes in the same manner as Fig. 3a except for the inclusion of the amount of FLD protein in the TTS region. FLD showed high importance. **(b)** ATXR7-bound TTS regions are highly enriched in FLD. **(c)** There is a significant overlap between genes that lose H3K4me1 in the atxr7 mutant (RPKM WT-atxr7 > 6.5) and genes that gain H3K4me1 in the fld mutant. p-value < 10e-6, hypergeometric test.

**Supplementary Fig. 8.**
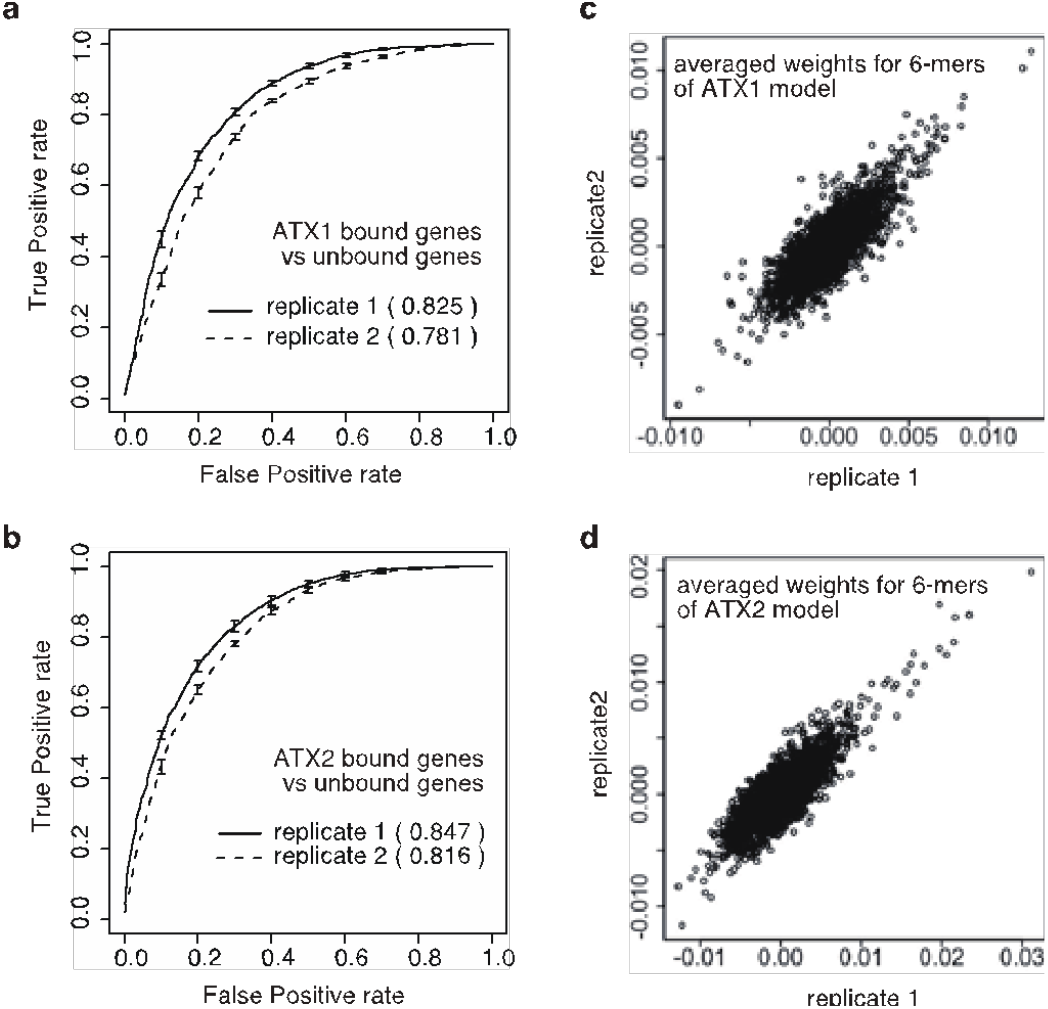
Biological replicates confirmed that DNA motifs also rule ATX1 and ATX2 localization. The ChIP-seq data sets of the biological replicates (Supplementary Fig. 6) were analyzed in the same manner as Fig. 4. Biological replicates also achieved accurate prediction for both **(a)** ATX1 and **(b)** ATX2 as demonstrated as ROC plot. **(c,d)** The learned trends of weights highly correlated between the replicates.

**Supplementary Fig. 9.**
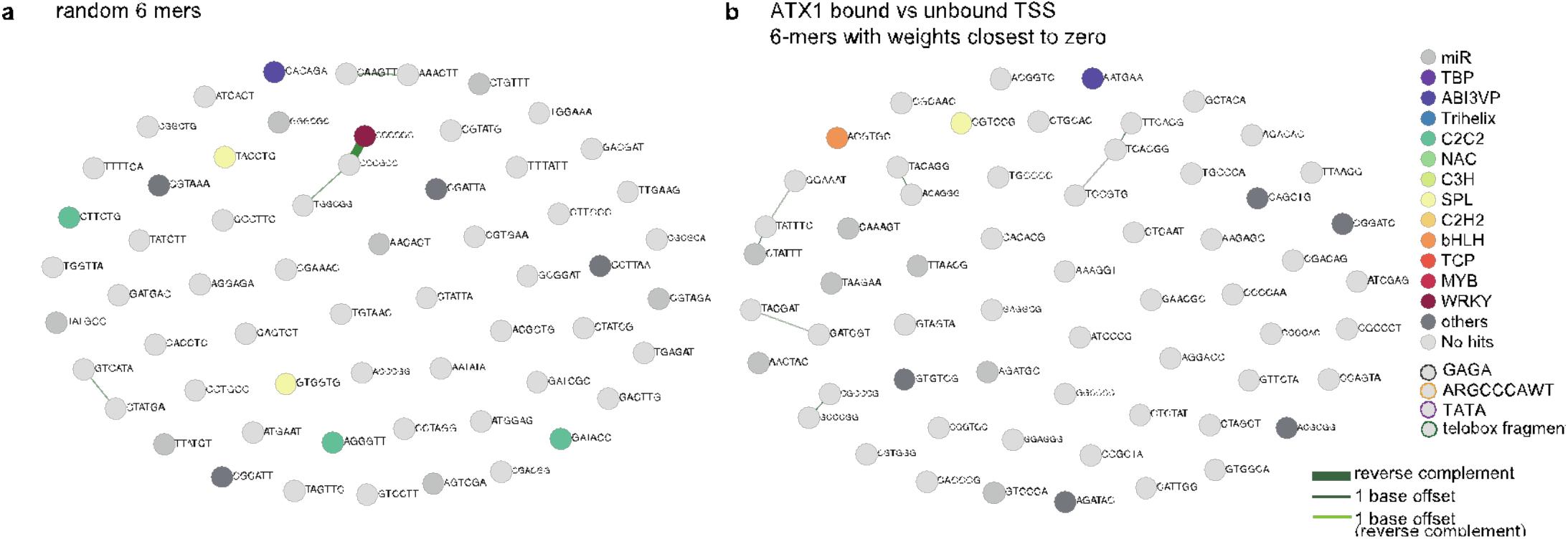
Non-predictive 6-mers. (a,b) Clustering and annotation of non-predictive sixty 6-mers. Random (**a**) and near-zero-weighted sixty 6-mers in the ATX1 model.

**Supplementary Fig. 10.**
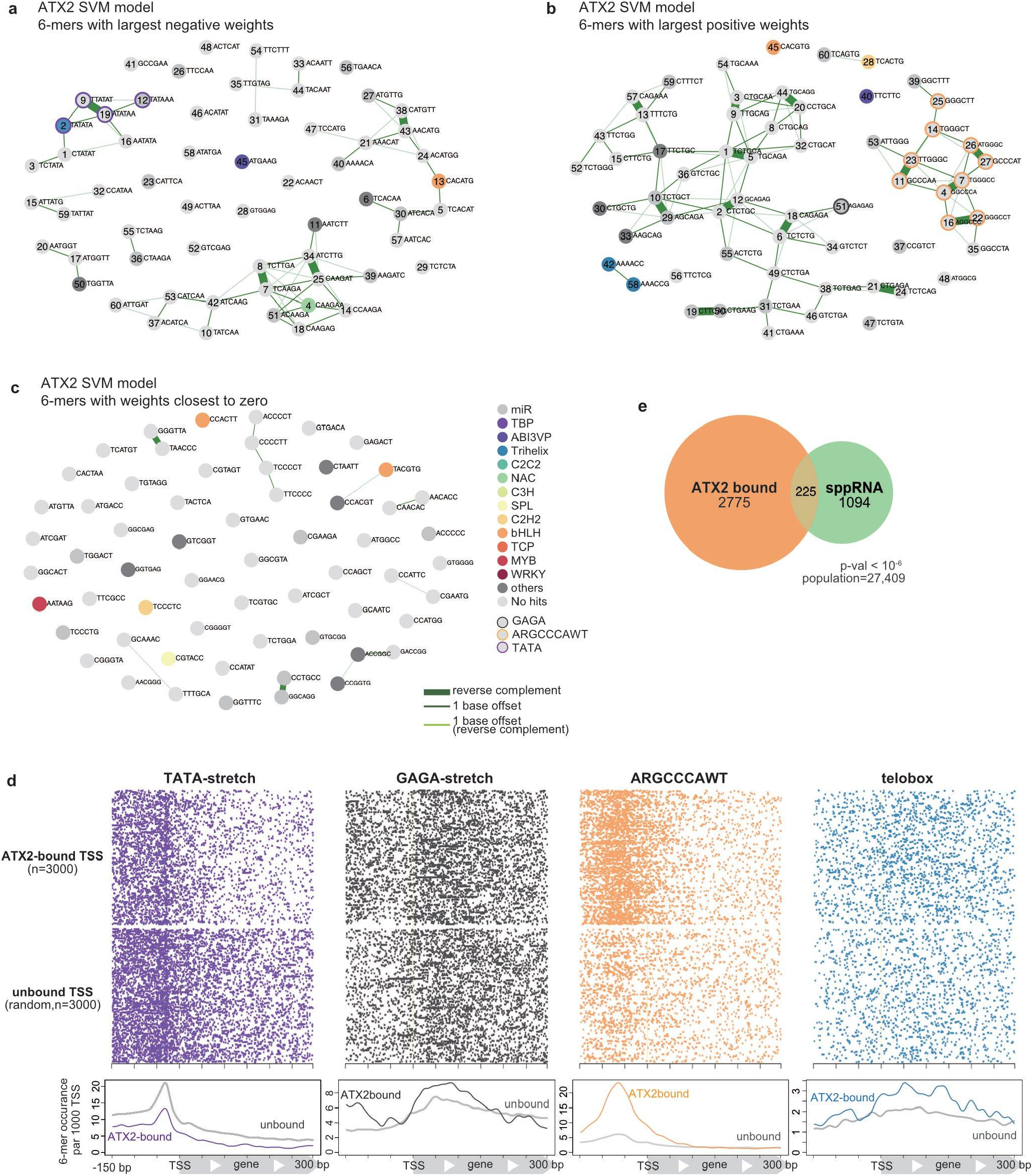
Annotations of predictive and non-predictive 6-mers in the ATX2 model. Predictive and non-predictive 6-mers in the ATX2 models were selected and analyzed in the same manner as in ATX1 models (Fig. 4). **(a-c)** Clustering and annotation of predictive and non-predictive 6-mers. Each circle represents the top sixty 6-mers with negative (**a**), positive (**b**), or nearly zero (**c**) SVM weights in the ATX2 model. **(d)** Positional distributions of highly weighted motifs in the TSS region. **(e)** ATX2-bound TSS significantly overlaps with sppRNA-harboring TSS detected in the *hen2-2* background. The significance of the overlap was tested using a hypergeometric test.

**Supplementary Fig. 11.**
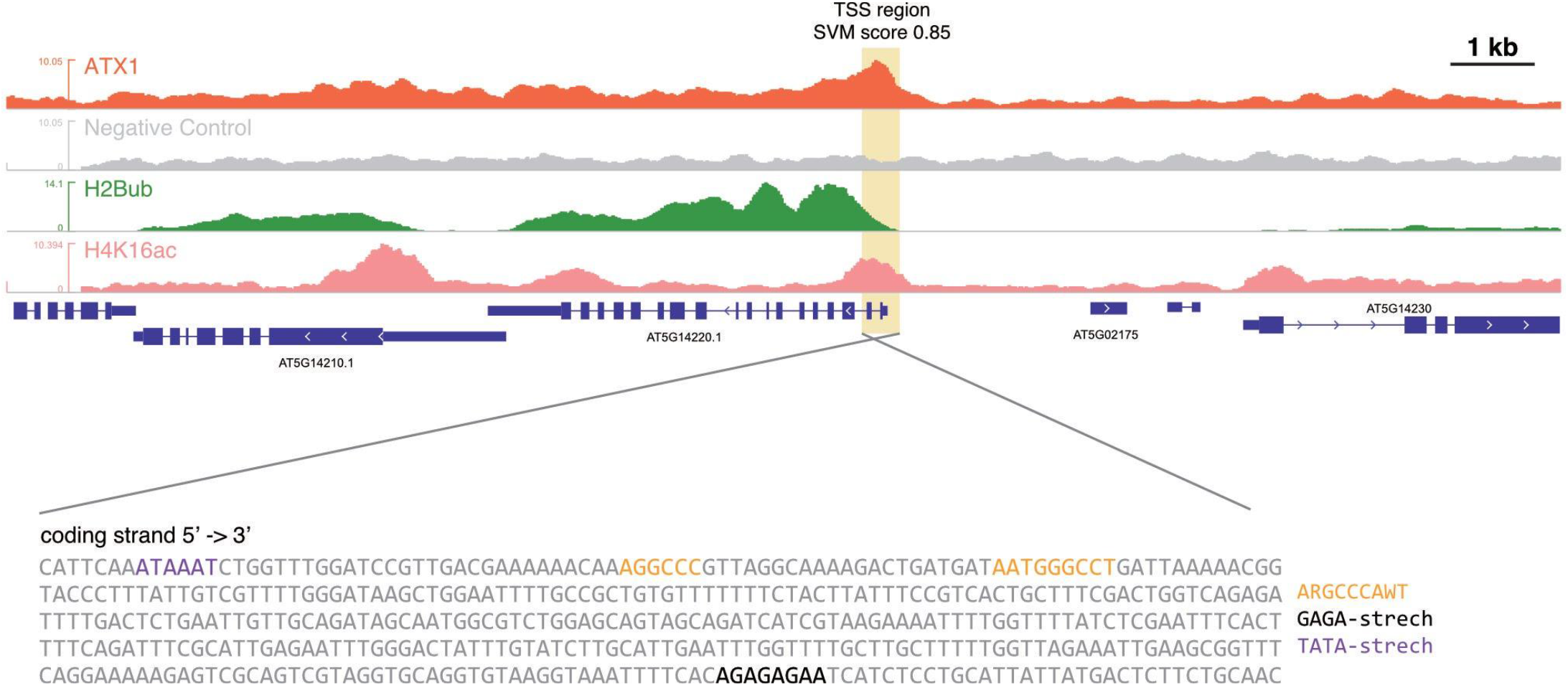
An example browser view of ATX1-bound and unbound genes. Examples of the ATX1-bound and unbound genes. ChIP-seq read coverage tracks of ATX1, negative control (non-transgenic), H2Bub, and H4K16c are shown. AT5G14220 is one of the ‘ATX1-bound genes’. Its ‘TSS region’, highlighted with yellow shade, has higher levels of H2Bub and H4K16ac compared to the other TSS regions in the view. The SVM score of this TSS region in the full model is 0.85 (ranges from 0 to 1. The higher the value is, the more likely it to be ATX1-bound, based on DNA sequence). In the DNA sequence of the TSS region, positively weighted ARGCCCAWT and GAGA-stretch are present, as well as negatively weighted TATA-stretch.

**Supplementary Fig. 12.**
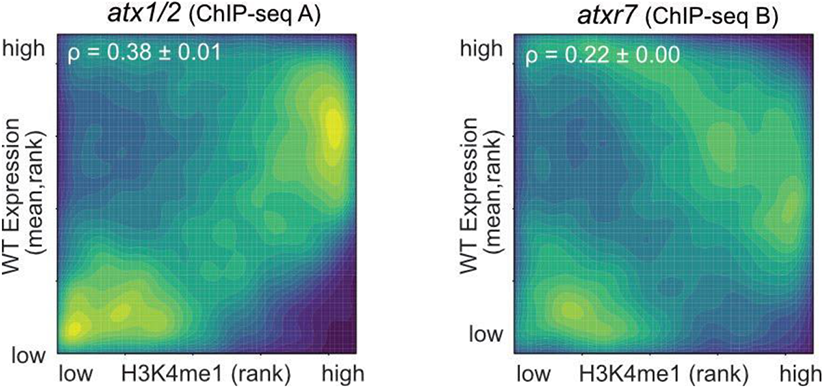
Change in H3K4me1-transcription correlation is due to change in H3K4me1, not in transcription. Correlation between H3K4me1 and expression is shown in the same way as Fig. 5, except that in this figure the expression (mRNA-seq) data are of WT. The *atx1/2* mutant has a higher, while *atxr7* has a lower correlation than WT, demonstrating that the change in H3K4me1 rather than transcription causes the change in the correlation. ρ is the correlation estimate of Spearman’s test. The color scale is the same as Fig. 5.

**Supplementary Fig. 13.**
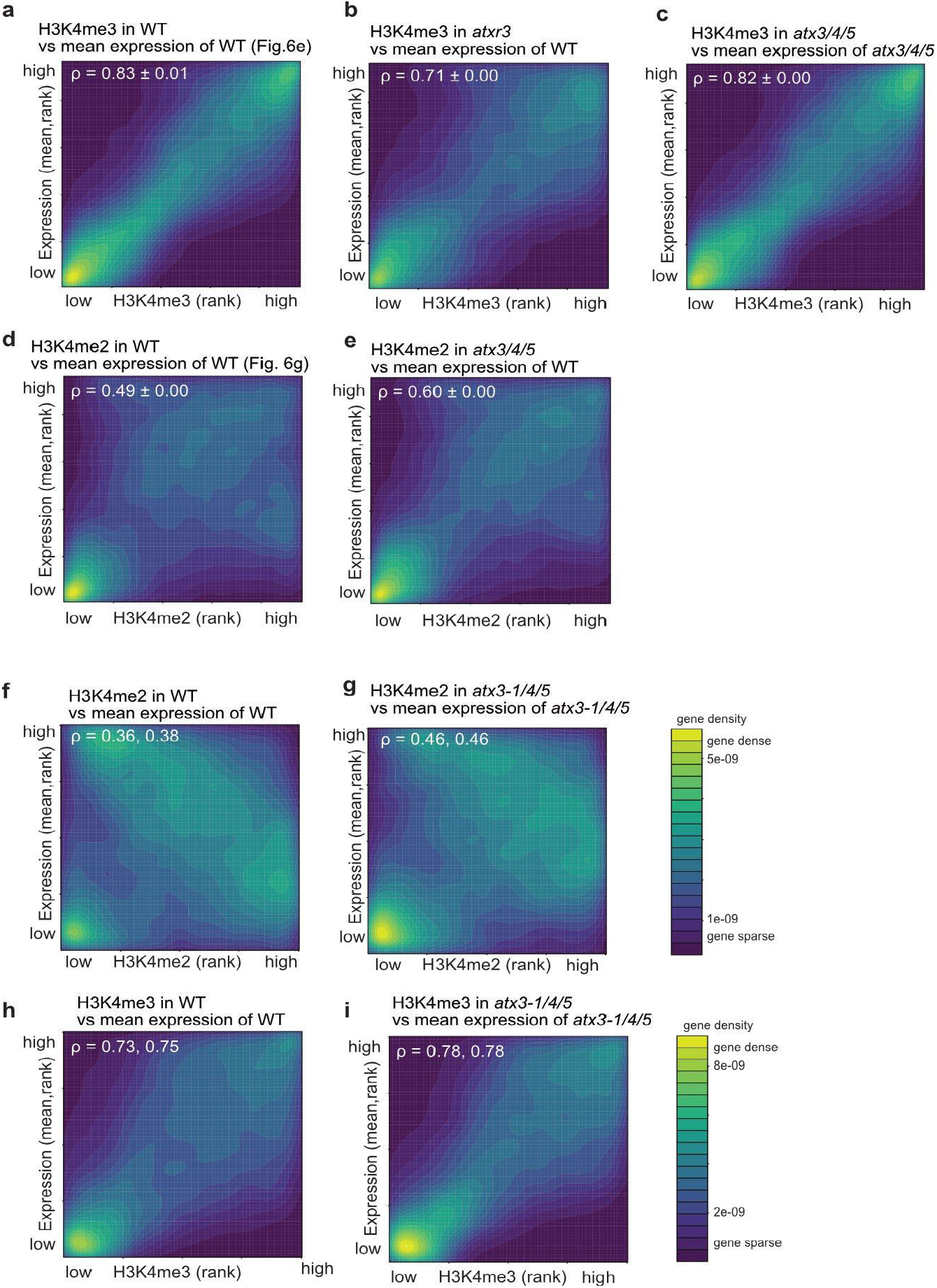
Correlational trend between H3K4me and transcription of *atxr3* and *atx3/4/5*. **(a)** Identical to Fig. 6e and shown for reference. **(b)** Correlation between H3K4me3 level in *atxr3* and expression level in WT. Compared to WT-WT comparison in **(a)**, the correlation is diminished and almost as weak as a mutant-mutant comparison in (Fig. 6f). **(c)** *atx3/4/5* mutant does not show a large correlational change between H3K4me3 and transcription. **(a-c)** and Fig. 6e,f are shown in the same color scale. **(d)** Identical to Fig. 6g. **(e)** Correlation between H3K4me2 level in *atx3/4/5* and expression level in WT. Compared to a WT-WT comparison in **(d)**, the correlation is stronger and almost as strong as a mutant-mutant comparison in Fig. 6h. **(f-i)** Change in the correlation between H3K4me3 **(f, g)** or H3K4me2 **(h, i)** and transcription in the *atx3-1/4/5* mutant, from Chen et al 2017’s datasets. *atx3-1/4/5* mutant, which has stronger effects on H3K4me2 and H3K4me3 compared with *atx3*/*4*/*5* (Fig. 1e, Supplementary Fig. 1), has a stronger correlation not only in transcription-H3K4me2 but also in transcription-H3K4me3.

**Supplementary Fig. 14.**
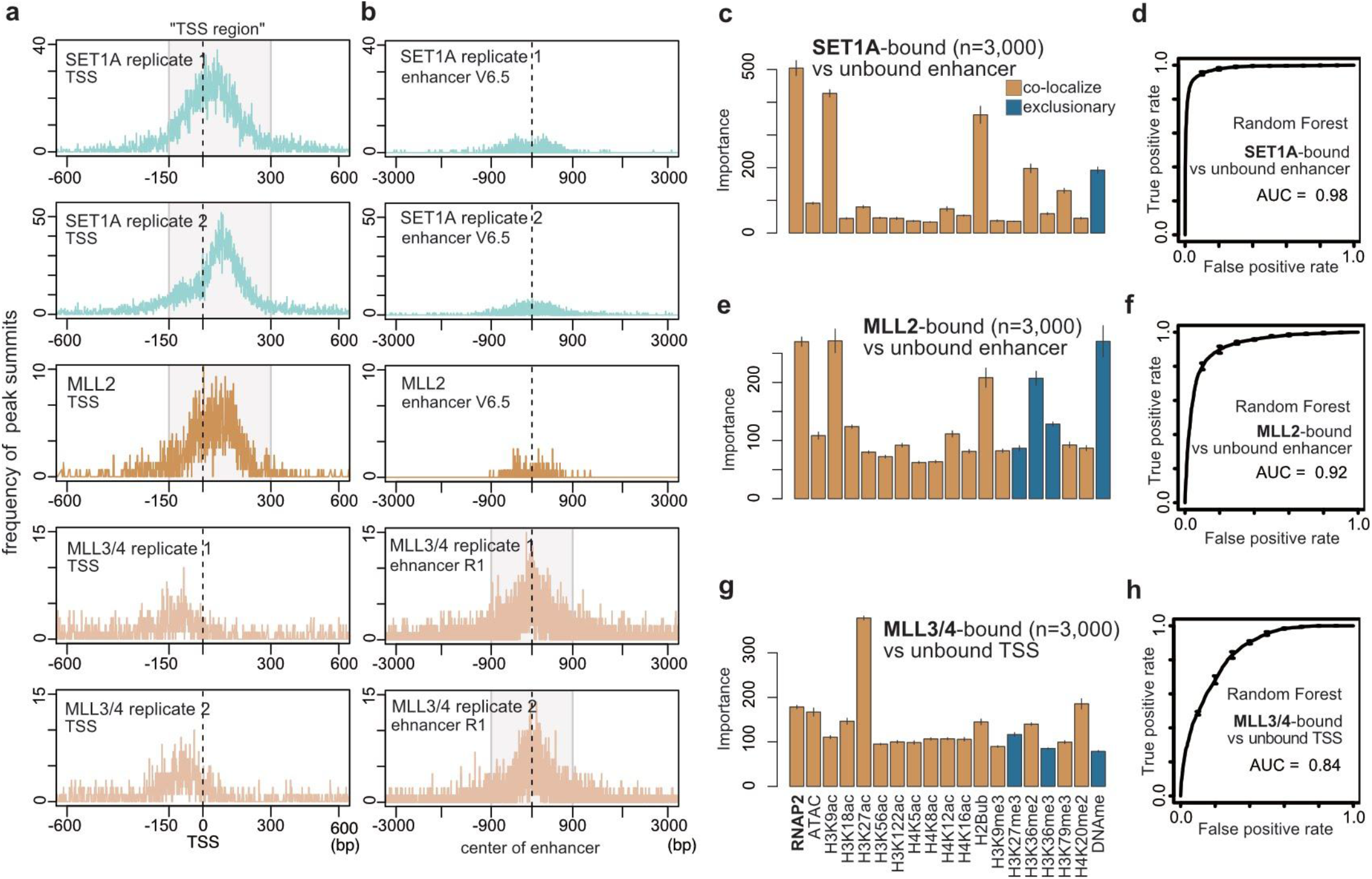
Localization of mammalian H3K4 methylases in TSSs and enhancers. **(a,b)** Position of mammalian H3K4 methylases’ peaks relative to TSS (**a**) and enhancer (**b**), visualized as a frequency of ChIP-seq peak summits. x-axis, distance from TSS (**a**) or center of enhancer (**b**); y-axis, number of peak summits. Most of SET1A and MLL2’s peaks belong to the region spanning from 150 bp upstream to 300 bp downstream of TSS, where we hereafter refer to as the ‘TSS region’. Most of the MLL3/4 peaks belong to 1800 bp around the center of the enhancer, where we hereafter refer to as ‘enhancer’. **(c,e,g)** ‘Importance’ of features derived from the Random Forest models trained to predict enhancer occupied by SET1A (**c**) and MLL2 (**e**), or to predict TSS region occupied by MLL3/4 (**g**). Graphs are shown like Fig. 7b,e,h. **(d,f,h)** ROC plot showing the prediction accuracy of the random forest models corresponding to (**c**,**e**,**g**). All ROC and AUC are calculated with test data (25% of the original data). Average and standard deviation of the 5 repeats of training are plotted.

**Supplementary Fig. 15.**
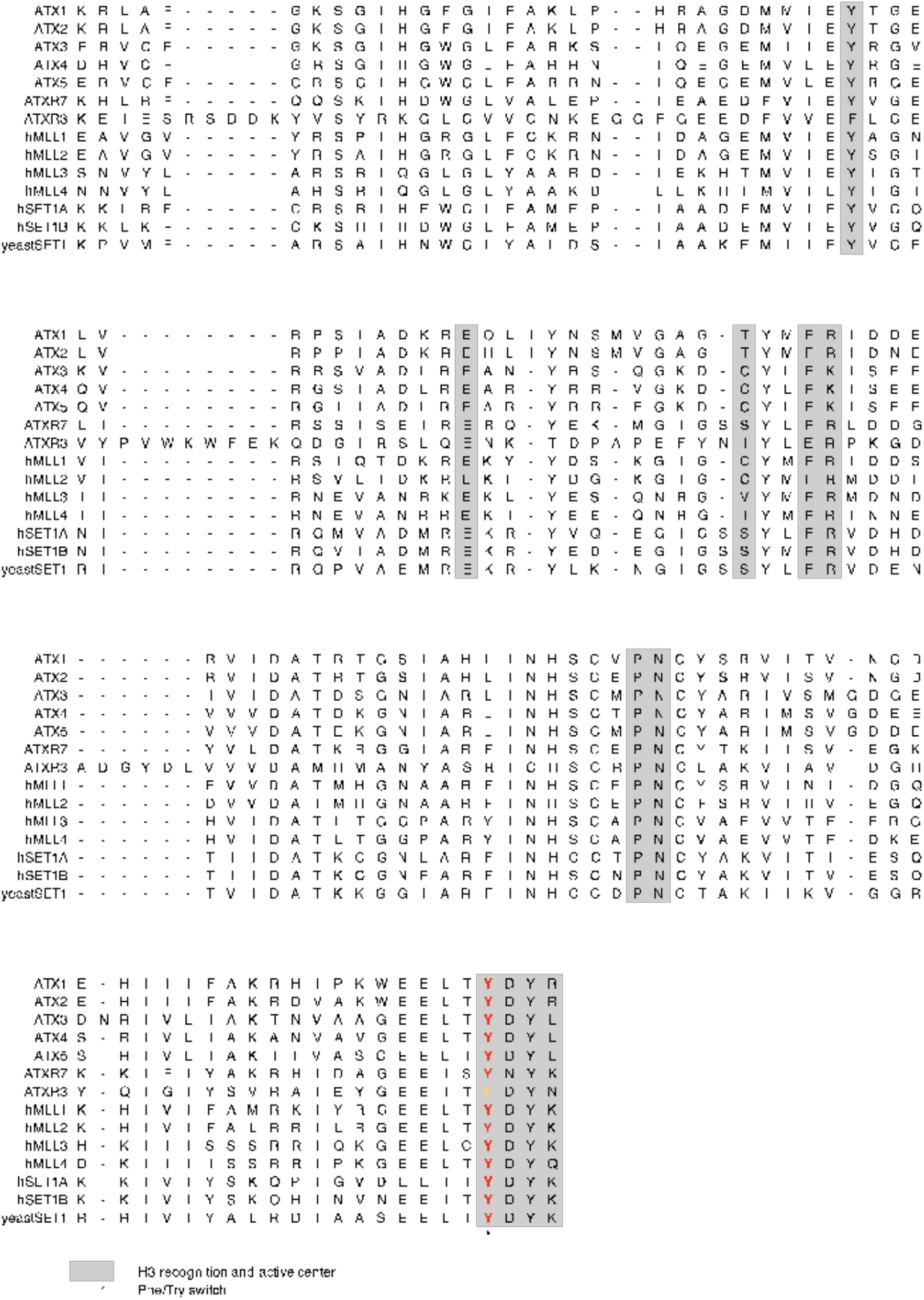
Amino acid alignment of catalytic domain SET in ATXs and their homologs. Y at ‘Y/F-switch’ is typical for SET domains of mono/di-KMTs ^44^. Annotations are from ref 91.

**Supplementary Fig. 16.**
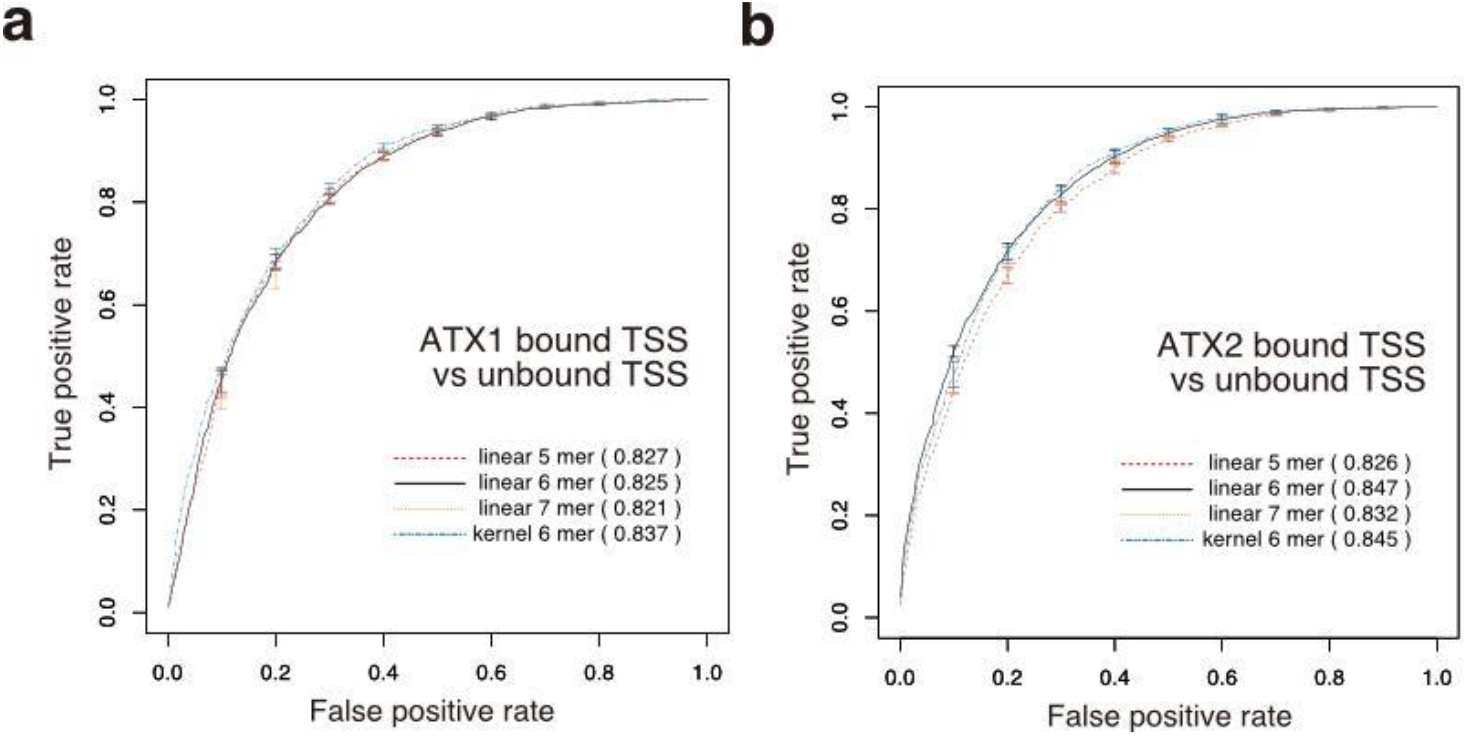
Parameter tuning of SVM models. **(a,b)** Prediction accuracy of linear SVM models trained with different sizes of k-mers, and kernel SVM with 6-mers. The numbers in parenthesis are AUC calculated with test data.

**Supplementary Table 1** List of all 6-mers and their weights in ATX1 and ATX2 models in replicates. Weights are averaged from 5 cross-validation models. ‘ATX1_model’ corresponds to a model in Fig. 4, and ‘ATX1_model_replicate’ corresponds to models based on biological replicate datasets shown in Supplementary Fig. 8.

**Supplementary Table 2** List of highly weighted and unweighted 6-mers, shown in Fig. 4e, f, Supplementary Fig. 9 and Supplementary Fig. 10a, b, c, and these 6-mer’s best TOMTOM hits that meet the criteria, q-value < 0.1.

**Supplementary Table 3** Sources of reanalyzed data in mESC.

## Notes

### Competing Interest Statement

The authors have declared no competing interest.

